# Somatic hypermutation analysis for improved identification of B cell clonal families from next-generation sequencing data

**DOI:** 10.1101/788620

**Authors:** Nima Nouri, Steven H. Kleinstein

## Abstract

**Motivation:** Adaptive immune receptor repertoire sequencing (AIRR-Seq) offers the possibility of identifying and tracking B cell clonal expansions during adaptive immune responses. Members of a B cell clone are descended from a common ancestor and share the same initial V(D)J rearrangement, but their B cell receptore (BCR) sequence may differ due to the accumulation of somatic hypermutations (SHMs). Clonal relationships are learned from AIRR-seq data by analyzing the BCR sequence, with the most common methods focused on the highly diverse junction region. However, clonally related cells often share SHMs which have been accumulated during affinity maturation. Here, we investigate whether shared SHMs in the V and J segments of the BCR can be leveraged along with the junction sequence to improve the ability to identify clonally related sequences. We develop independent distance functions that capture junction similarity and shared mutations, and combine these in a spectral clustering framework to infer the BCR clonal relationships. Using both simulated and experimental data, we show that this model improves both the sensitivity and specificity for identifying B cell clones.

**Availability:** Source code for this method is freely available in the **SCOPer** (Spectral Clustering for clOne Partitioning) R package (version 0.2 or later) in the Immcantation framework: www.immcantation.org under the CC BY-SA 4.0 license.

## 1 Introduction

B cells recognize pathogens through their B cell receptor (BCR). The ability to recognize and initiate a response to a wide variety of pathogens depends upon a large population of B cell lymphocytes each of which expresses a particular receptor for antigen. The diversity of the BCRs (also referred to as Immunoglobulin (Ig) receptors) is a result of genetic recombination and diversification mechanisms. BCRs are comprised of two identical heavy (IGH) and light (IGL) chain proteins. For IGH-chains, diversity is initially created in the germline via recombination of variable IGHV,diversity IGHD, and joining IGHJ genes (termed the V(D)J recombination process (Tonegawa, 1983)). Diversity in IGH is further increased by addition of P- and N-nucleotides at the IGHV/IGHD and IGHD/IGHJ boundaries (Alt and Baltimore, 1982; Lafaille *et al.*, 1989; Murphy, 2011). For IGL-chains, the IGLV gene is rearranged directly to IGLJ gene. The region where IGHV, IGHD and IGHJ come together in IGH (or IGLV and IGLJ for IGL) is termed the CDR3 (the junction region is defined as the CDR3 plus the prefix and suffix conserved flanking amino acid residues), and this high diversity region is often involved in antigen-binding (Xu and Davis, 2000).

During T-dependent responses, antigen-activated B cells undergo clonal expansion and acquire additional diversity through somatic hypermutation (SHM), an enzymatically-driven process introducing point substitutions into the BCR locus at a rate of ∼ 1/1000 bp/cell division (McKean *et al.*, 1984). B cells that acquire mutations that improve their ability to bind the pathogen are preferentially expanded leading to affinity maturation of the B cell population over time. Therefore, SHMs have important consequences for the kinetics, quality, and magnitude of B cell clones as the fundamental building blocks of immune repertoires (Kepler and Perelson, 1993).

Accurate identification of clonal relationships is important, as these clonal families form the basis for a wide range of repertoire analyses, including diversity analysis (Robins *et al.*, 2013; Meng *et al.*, 2017; Rosenfeld *et al.*, 2018), lineage reconstruction and detection of antigen-specific sequences (Yaari and Kleinstein, 2015; Tsioris *et al.*, 2015; Hoehn *et al.*, 2019) and effector functionality (McKean *et al.*, 1984; Sablitzky *et al.*, 1985). One way to monitor and track B cell clonal lineages is to perform large-scale sampling of B cell populations, amplifying, and sequencing the expressed antibody gene rearrangements by next-generation sequencing (NGS) (Ansorge, 2009; Weinstein *et al.*, 2009; Boyd *et al.*, 2009; Metzker, 2010). Recent studies by NGS have greatly expanded our understanding of B cell clonal lineage development in high-throughput Adaptive Immune Receptor Repertoire sequencing (AIRR-seq) data (Boyd and Joshi, 2015; Rubelt *et al.*, 2017; Vander Heiden *et al.*, 2018). However, clonal relationships are not directly measured, but they must be computationally inferred. To this end several computational methods have been proposed to identify B cell clones from high-throughput AIRR-seq data (Glanville *et al.*, 2011; Kepler, 2013; Ralph and Matsen IV, 2016; Gupta *et al.*, 2017; Nouri and Kleinstein, 2018b).

Antibody diversity is largely dominated by the IGH-chain (Xu and Davis, 2000). The IGH-chain owes this diversity to the: (1) use of an IGHD gene, which IGL-chains lack, (2) addition of short palindromic (P) nucleotides at the IGHV-IGHD and IGHD-IGHJ joints (Lafaille *et al.*, 1989), (3) insertion of non-templated (N) nucleotides at the IGHV-IGHD and IGHD-IGHJ joints by terminal deoxynucleotidyl transferase (TdT) (Alt and Baltimore, 1982), and (4) higher rates of SHM than IGL-chains (Wood *et al.*, 2001). The IGH-chain junction region commonly serves as an identifier for clonal inference methodologies. For instance, sequences whose junctions are identical or have a high degree of homology (measured by string distance at the nucleotide level) are often classified as belonging to the same clone (Hershberg and Prak, 2015). However, to avoid grouping together highly homologous yet distinct sequences, some studies also regroup sequences to have the same IGHV- and IGHJ-gene annotations to be considered clonally-related (Zhang *et al.*, 2015). Many methods also assume that members of a clone share the same junction length, because SHMs introduced into the BCR sequence are predominantly point substitutions. Probabilistic models have also been developed to calculate the likelihood of sharing a common B cell ancestor and subsequently infer clonal grouping (Kepler, 2013; Ralph and Matsen IV, 2016). However, these methodologies have complexities that become substantially expensive for large sequencing datasets. Overall, in practice, a common approach is to infer clones among sequences with high junction region similarity, as well as identical junction length and IGHV- and IGHJ-gene usage (referred to as recombination-based model) (Hershberg and Prak, 2015).

While recombination-based strategies are common among current studies, clonal relationship inference solely based on the similarity of the junction region does not leverage the potential information in the V and J segments. It has been suggested that incorporating shared SHMs in these regions could improve recombination-based clonal inference (Zhou and Kleinstein, 2019). Members of an expanded B cell clone often share specific somatic mutations and, sometimes, combinations of mutations across the BCR. Mutations may be shared among two or more members of a clone as a simple result of being passed down during cell division, or may be positively selected as part of the affinity maturation process (Clarke *et al.*, 1985; Blier and Bothwell, 1987; Diamond *et al.*, 1992; Coker *et al.*, 2003; Furuta *et al.*, 2017). This hierarchy of shared mutations can be considered as the “glue” binding all the members of a B cell clone together and shaping its lineage tree (Figure 1). This additional IGH-chain information could be leveraged to refine clonal relationships.

**Fig. 1.**
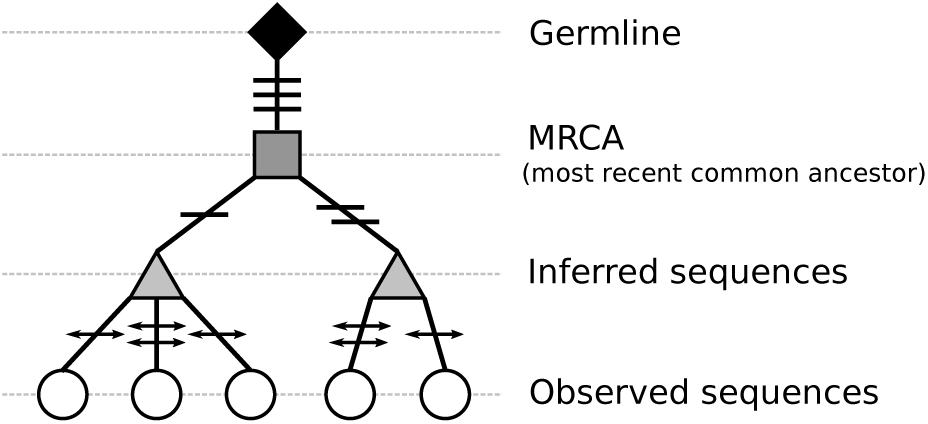
A B cell lineage tree showing the relationships between clonally-related cells. The germline sequence (diamond) is shown at the root of lineage, and is connected by a single branch to the most recent common ancestor (MRCA) (square). This branch consists of mutations that are shared across all members of a clone. Several sub-branches descend from the MRCA to inferred sequences (triangles) carrying mutations that are shared by a subset of clone members. Finally, the inferred sequences are connected to observed sequences (circles) through mutations that are unique to each given observed sequence. Shared and unique mutations are marked at each branch by horizontal lines and arrowhead-lines, respectively.

In this study, we investigated whether shared SHM patterns in the V and J segments of the BCR can be leveraged along with the junction sequence to improve the ability to identify clonally related sequences. This model is implemented in the new version of **SCOPer**. The first version of **SCOPer**, a spectral clustering-based method for identifying clones from high-throughput B cell repertoire sequencing data, was presented in Nouri and Kleinstein (2018b). In the following sections, we discuss the main steps of the methodology and explain our implementation of the recent improvements upon the original framework. We further examine the performance of **SCOPer** using simulated and experimental datasets.

## 2 Method

The clonal inference procedure by **SCOPer** is composed of four main steps (Figure 2). First (1), BCR sequences IGHV and IGHJ genes are identified. This can be done using various publicly available tools such as IMGT/HighV-QUEST (Alamyar *et al.*, 2012) or IgBLAST (Ye *et al.*, 2013). Then (2), sequences are partitioned into groups (termed as “VJ(*ℓ*)-group”) that share the same IGHV- and IGHJ-gene (gene-level grouping) and junction length (length-level grouping). The gene-level grouping is based on the assumption that the identity of germline gene (the clone members unmutated common ancestor) cannot change through affinity maturation. The length-level grouping is based on the assumption that sequences evolve only through point mutation (no indels). Next (3), within each given VJ(*ℓ*)-group the defining metric that indicates common clonality among BCR sequence pairs is determined by combining the similarity among junction region sequences (recombination-based distance: subsection 2.1) and the V and J segment mutation profile (SHM-based distance: subsection 2.2) into an integrated distance function (subsection 2.3). Finally (4), BCR clones are identified using spectral clustering-based approach built upon this distance function (subsection 2.4).

**Fig. 2.**
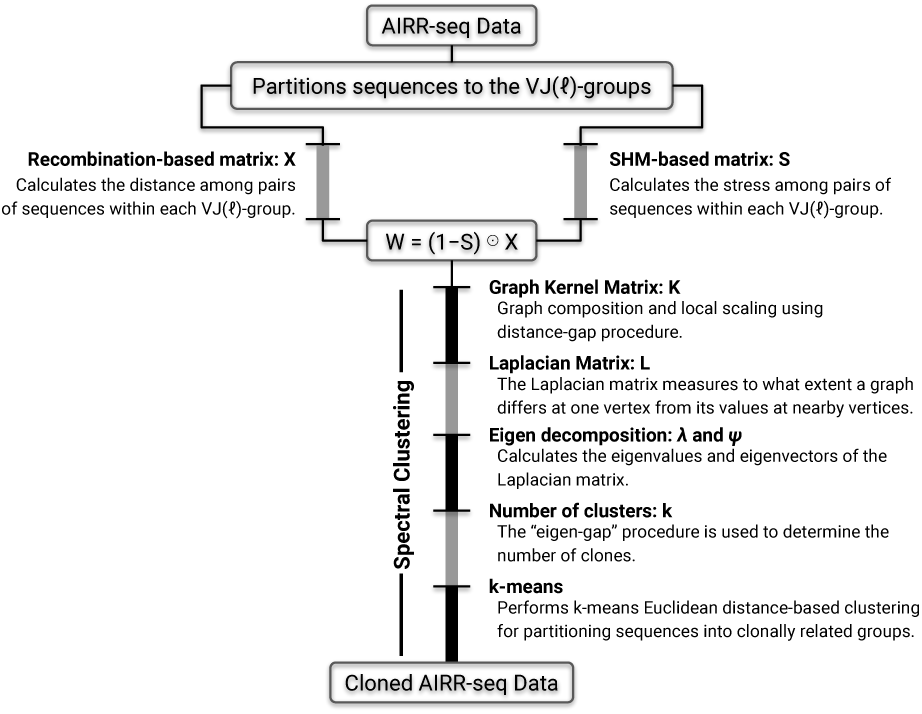
Overview of the **SCOPer** workflow. First, AIRR-seq data are partitioned into VJ(*ℓ*)-groups which contain sequences with the same IGHV gene annotation, IGHJ gene annotation, and junction length. Next, each VJ(*ℓ*)-group is subject to a recombination-based and a SHM-based distance calculation. Finally, the outputs of these calculations are combined into an integrated distance function that is used as the basis for inferring the BCR clonal relationships using a spectral clustering-based approach.

### 2.1 Recombination-based distance matrix calculation

The recombination-based component of **SCOPer** is focused on the sequencing reads’ junction region. At this step, we generate a symmetric and positive pair-wise similarity matrix *X*_*ij*_ defined by the Hamming distance between the junction regions corresponding to the *i*^th^ and *j*^th^ sequences from a given VJ(*ℓ*)-group. This is called the “junction-targeted” recombination-based distance matrix. The Hamming distance is defined as the number of positions at which the corresponding nucleotides are different. The recombination-based distance matrix can also be generated from CDR3 region by excluding the three-nucleotide prefix and suffix from both ends of the junction (i.e. converting junction segment to CDR3 region). Henceforth, this is called a “CDR3-targeted” recombination-based distance matrix.

### 2.2 SHM-based distance matrix calculation

The SHM-based component of **SCOPer** is focused on the V and J segments. We develop a model in which the occurrence of a mutation at the same nucleotide position of a pair of sequences (referred to as “pair-wise shared mutation”) will be used, accompanying with recombination-based component, in order to define a metric that indicates common clonality among BCR sequence pairs. We generate a SHM-based distance matrix so that a pair of sequences with a higher shared mutation rate are more likely to belong to the same clone, whereas a pair of sequences with a lower shared mutation rate are considered more independent from each other. By analogy with continuum mechanics, the pair-wise shared mutations can be loosely referred to as “stress” which expresses the internal forces that clonally related sequences exert on each other.

We begin with identification of the pair-wise mutations. First, for each VJ(*ℓ*)-group a single germline representative is generated by building the effective sequence of all germlines (allele-grouping). This will facilitate identification of the pair-wise mutations in VJ(*ℓ*)-groups whose germlines have different nucleotides at the same position (alleles). The representative germline is deterministic such that if a position contains different nucleotides, the effective will be an IUPAC (International Union of Pure and Applied Chemistry) character representing all of the nucleotides present. Henceforth, we refer to such sequence as “effective germline”. Then, in each VJ(*ℓ*)-group, pairs of sequences are compared with the effective germline to identify mutations. We note that, depends upon the type of recombination-based matrix calculated in the previous step (i.e., junction- or CDR3-targeted) the junction or CDR3 region of the sequences and germlines are excluded from this analysis.

We continue with a categorical approach to classify the identified pair-wise mutations (Figure 3). For each pair of *i*^th^ and *j*^th^ sequences the mutations at each position are flagged with a binary variable and categorized in three classes: (1) a single mutation which occurs only in one of the sequences, 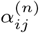, (2) two unique mutations which occur in both sequences, 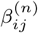, and (3) a shared mutation which occurs in both sequences, 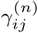. Here, the parameter *n* indicates the position of each nucleotide along the sequence string. The binary variables are retrieved to create two matrices. One of the matrices accumulates the total number of mutations:

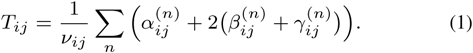

A second matrix accumulates the shared mutations:

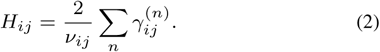

Here, *T*_*ij*_ is a positive value and always larger than or equal to positive value *H*_*ij*_. The term *ν*_*ij*_, average number of informative positions (*∈* {A,C,G,T}) in *i*^th^ and *j*^th^ sequences, is a normalizing factor used to prevent bias toward pairs of sequences with fewer non-ACGT positions.

**Fig. 3.**
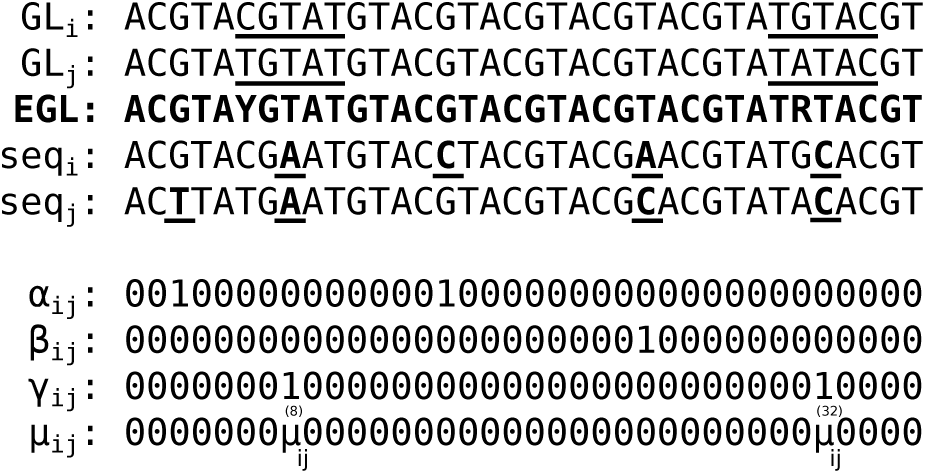
Pair of sequences (seq) are compared with each other and the VJ(*ℓ*)-group effective germline (EGL) to identify unique and shared somatic hypermutation events. The effective germline sequence is determined by IUPAC character representation of all the nucleotides present at each position across all germlines in a given VJ(*ℓ*)-group (allele-grouping). Each nucleotide position of *i*^th^ and *j*^th^ sequences is compared with the corresponding nucleotide position in the effective germline and somatic hypermutation events are flagged with binary variables: (1) *α*: a single mutation which occurs only in one of the sequences, (2) *β*: two unique mutations which occur in both sequences, and (3) *γ*: a shared mutation which occurs in both sequences. The average of the mutabilities of the germlines (GLs) 5-mer motifs in which a shared mutation occurred at the central position is shown by 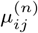, where superscript *n* indicates the position that mutation occurred. Mutation events are bold and underlined in the sequences.

We note that SHM biases have been reported (Elhanati *et al.*, 2015; Yeap *et al.*, 2015) both in the bases that are targeted (Betz *et al.*, 1993; Shapiro *et al.*, 2003) as well as the substitutions that are introduced (Smith *et al.*, 1996; Cowell and Kepler, 2000). These biases have been summarized by hot- and cold-spot targeting model (“S5F” model that produces background likelihood of a particular mutation based on the surrounding sequence context as well as the mutation itself) by *Yaari et al.* (2013). We reasoned that mutations at hot-spot positions could be more likely to be shared by sequences that are not truly clonally related. In order to account for the potential influence of SHM biases, we incorporate a damping matrix in the form of:

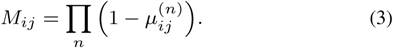

Here, 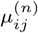 is the average of the mutabilities of the germlines micro-sequence motifs (e.g., a 5-mer from “S5F” model) in which a mutation occurs at the central position *n*. Each value is subtracted from one to reverse the scaling direction (Figure 3).

We finalize the SHM-based step by calling equations 1, 2, and 3 to calculate the stress between *i*^th^ and *j*^th^ sequences in the form of:

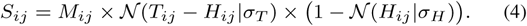

Here, 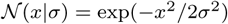 is a continuous Gaussian probability distribution, where parameter *σ_T_* and *σ_H_* are the standard deviations of the *T* and *H* matrices capturing the variability of total and shared SHM events in each VJ(*ℓ*)-group, respectively. It is important to note that for different VJ(*ℓ*)-groups the level of similarity that indicates common clonality may be different. Therefore, using the Gaussian probability distribution, built upon the given VJ(*ℓ*)-group, will make the model capable of adapting itself to the local mutation frequency.We further note that, the stress becomes nonzero only if the number of pair-wise shared mutations is non-zero (*H* ≠ 0). Conversely (i.e., *H* = 0), the stress is forced to zero by the third term of Eq. 4, even though non-shared mutations exist (*T* ≠ 0), and consequently the recombination-based part of the **SCOPer** is fully in charge to infer the clonal relationships. In practice, the behavior of the stress function (Eq. 4, ignoring the impact of SHM hot-spots, i.e. *M*_*ij*_ = **1**) comparing two pairs of sequences can be described as follows:

- if no shared mutations are observed, then the stress *S*_*ij*_ is zero,
- if the two pairs have the same total number of mutations, then the pair which accumulates more shared mutations will have higher stress, and
- if the two pairs have the same number of shared mutations, then the pair which accumulates fewer non-shared mutations, will have higher stress. (Note that *T*_*ij*_ is always larger than or equal to *H*_*ij*_).

### 2.3 Graph composition and local scaling

The spectral clustering at the core of **SCOPer** works based on a graph construction procedure where the vertices are the observed sequences to be clustered, and the edges between vertices are weighted dependencies among pairs of sequences. The graph construction relies on a quantitative notion of adaptive local neighborhoods in the dataset, which are encoded by a symmetric Kernel function. The Kernel function is used to capture intrinsic data geometries that approximate underlying manifold models from the data. To construct the kernel graph, first, we generate a weighted-distance matrix in the form of,

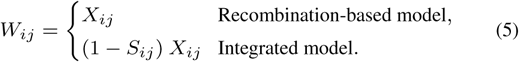

The model is named “recombination-based” when recombination-based distance matrix (*X* from subsection 2.1) is only involved in graph composition. The model is named “integrated” when recombination-based (*X* from subsection 2.1) and SHM-based (*S* from subsection 2.2) distance matrices are both involved in graph composition. In integrated model, each stress value *S*_*ij*_ is subtracted from one to reverse the scaling direction, so that the pair of sequences (i.e., the graph vertices) with higher stress become closer to each other, thereby more likely to belong to the same clone. The integrated model can be loosely thought of as Hooke’s Law (*W* = *κX*, where *κ* = 1 *− S*), which rules the attraction force between a pair of sequences using a “spring” with proportionality factor *κ* (see Figure 4). In the subsequent step, we generate a fully connected graph Kernel using a Gaussian similarity function in the form of,

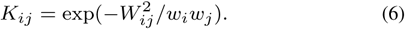

**Fig. 4.**
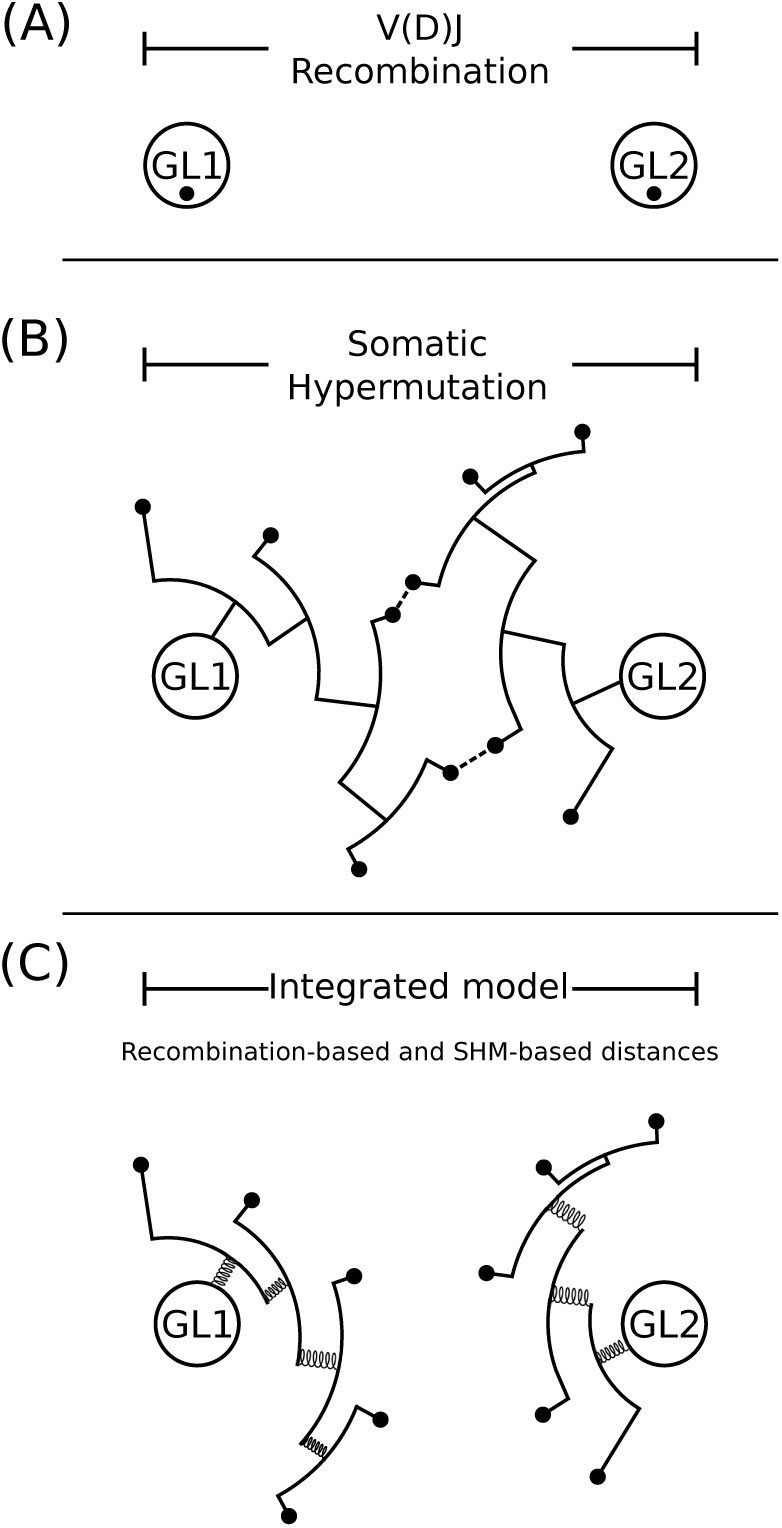
The integrated model pulls together clonally-related sequences to improve the B cell clonal inference process. (A) V(D)J recombination generates a set of highly diverse (unmutated) sequences with large distances between independent clones (inter-clonal diversity). (B) Clonal expansion with SHM adds additional diversity, and leads the sequences to spread out around the initial points of creation (intra-clonal diversity). Some sequences from independent clones could end up with CDR3s that start to look similar (dashed-lines), and may lead to false positives in the clonal relationship inference process. (C) The stress between pairs of sequences, expressed via shared mutations, acts as a spring that pulls clonally-related sequences toward each other resulting in a more accurate distinction of local neighborhoods. Black circles indicate observed sequences, while white circles indicate germlines (GL1 and GL2).

Here, parameters *w*_*i*_ and *w*_*j*_ are the scaling distances corresponding to the *i*^th^ and *j*^th^ sequences, respectively, which control the width of local neighborhoods allowing the level of similarity to vary in different parts of the graph. In this way, the local neighborhoods are determined for each sequence, instead of selecting an universal scaling parameter for all. The width of each local neighborhood is identified by a single weighted-distance value such that sequences inside the neighborhood are more similar to each other than the outsider sequences. In order to determine the sequence-to-sequence scaling parameters a self-tuning framework (Zelnik-Manor and Perona, 2005) (the so-called distance-gap procedure) is incorporated into **SCOPer**. The distance-gap procedure determines the scale parameter *w*_*i*_ corresponding to the *i*^th^ sequence by seeking a relatively large gap in the set of weighted-distances from *i*^th^ sequence to the rest of the sequences. The distance-gap pipeline is performed as follows. First, the set of weighted-distances corresponding to the *i*^th^ row of the matrix *W* is retrieved. Then, a binned Gaussian kernel density estimate of the weighted-distances is generated using the `density` function from the **stats** R package. Next, the set of extrema of the continuous density distribution is flagged by finding the weighted-distances at which the first derivative of the distribution is zero while the second derivative is positive, indicating a local minimum following a local maximum. Recall from univariate Calculus that the first and second derivative for some function *f* (*x*) corresponds to the slope of the tangent line and curvature of *f* at point *x*, respectively. Finally, the scale parameter *w*_*i*_ associated with *i*^th^ sequence is determined as the closest smaller weighted-distance to the extremum with the lowest density value. If such an extremum is not found, the scale parameter *w*_*i*_ is simply determined as the first largest gap of the rank-ordered set of entries corresponding to the *i*^th^ row of the matrix *W*.

Local scaling is especially useful when the classification of the B cell repertoire contains multiple scales (e.g., if one clone is tight, while another one is sparse). By means of local scaling, the junction sequence similarities between different clones are lower than the similarities within any single clone. Therefore, edges between sequences in local neighborhoods are connected with relatively high kernels (i.e., *K*_*ij*_ → 1), while edges between far away sequences have smaller kernels (i.e., *K*_*ij*_ → 0). This is an important advantage of this methodology, by allowing the level of sequence similarity to vary in different local neighborhoods (a biologically plausible assumption), over other methodologies that partition sequences using an universal (fixed) level of similarity overall the sequences (Gupta *et al.*, 2017).

### 2.4 Spectral decomposition and clustering

Having defined a scheme to set the graph scale parameters automatically, following with the calculation of the graph Kernel matrix *K*, the last unknown free parameter in the model is the number of clones *k*, which is determined by the eigen-decomposition of the Laplacian matrix. First, the Laplacian matrix *L* = *D − K* is calculated, where *D* is the diagonal matrix with its *i*^th^ diagonal element being the sum of *i*^th^ row of *K*. Then, the Laplacian matrix is eigen-decomposed with eigenvalues {0 = *λ*_1_ ≤ *λ*_2_ ≤ … ≤ *λ*_*m*_} and corresponding eigenvectors 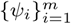, where *m* indicates the number of sequences. Then, the number of clones *k* is determined by finding the largest gap within the eigenvalue spectrum (the so-called “eigen-gap” procedure) at which adding another clone does not give much better modeling of the data. Finally, we perform *k*-means Euclidean distance-based clustering over the *k* eigenvectors 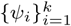 associated with the smallest *k* eigenvalues to find the members of each clone.

## 3 Bulk B cell simulation and library preparation

Each simulated dataset was generated using the **AbSim** R package (version 0.2.6) in a B cell single-lineage fashion (Yermanos *et al.*, 2017). Each B cell clone simulation begins with a random selection from sets of IGHV, IGHD, and IGHJ germline sequences (Giudicelli *et al.*, 2004) to produce a unique V(D)J recombination event. Then, clones are made by introducing mutations using a local nucleotide context-dependent model (i.e., S5F model from Yaari *et al.* (2013)), along a phylogenetic tree in which branching events occur stochastically. This process was repeated to create a collection of 25 simulated datasets. The size of each repertoire was sampled from a normal distribution (mean equal to 600k and standard deviation equal to 100k) and the clone sizes were sampled from a gamma distribution (shape equal to 0.75, scale equal to 0.75, and amplitude sampled from a normal distribution with mean equal to 1k and standard deviation equal to 0.1k). The remaining parameters were set as default. After simulation was done, the V and J annotations along with the junction segment of each simulated sequence were identified using IgBLAST version 1.13.0 (Ye *et al.*, 2013). Then, the outputs were retrieved and tab-delimited database files were generated using the command line tool `MakeDb`, from **Change-O** (version 0.4.5) (Gupta *et al.*, 2015). Quality checks were also undertaken to remove non-productive sequences. Specifically, each sequence was checked to satisfy a set of constraints that the: (1) whole sequence be annotated as functional, (2) whole sequence contains no stop codons, and (3) junction is in-frame (i.e. the length is modulo 3). Sequences which did not meet these criteria were excluded. At this point, sequences that are identical (i.e. copies that were generated coincidentally) are grouped together into “unique sequences”. The simulated datasets were further processed using the **SHazaM** (version 0.1.11 or newer) and **Alakazam** (version 0.2.11 or newer) R packages from Immcantation framework (www.immcantation.org) resulting in new columns containing VJ(*ℓ*)-group identifiers, mutation frequencies, and distance-to-nearest values (i.e., distribution of normalized Hamming distances from each junction sequence to its nearest non-identical neighbor in a given VJ(*ℓ*)-group). Finally, the outcome was a single tab-delimited file per each simulated dataset containing the metadata information associated with each sequence to be used as input to the clonal inference pipeline.

Table 1 presents an overview of 25 simulated datasets used in this study. Furthermore, the global metrics of the BCR simulated repertoires, including: (1) junction length distribution, (2) distance-to-nearest distribution, (3) clonal relative abundance distribution, (4) clone size distribution, (5) mutation frequency distribution, (6) number of clones per VJ(*ℓ*)-group, (7) average pair-wise SHM for clone, and (8) negative-control test, are presented in Supplementary Figures 1-25A-H, respectively.

**Table 1.** Overview of 25 simulated datasets generated by the **AbSim** R package (Yermanos et al., 2017). Each B cell clone is generated by one set of randomly selected unmutated human IGH-chain germline gene sequences (Giudicelli et al., 2004) to produce the V(D)J recombination event. Then, the germline undergoes clonal expansion along a phylogenetic tree in which branching events occur stochastically. SHM along this tree is modeled using a local sequence context-dependent model (i.e., “S5F” model from (Yaari et al., 2013)).

| Simulation | Sequences | Clones | Unique V-genes | Unique J-genes |
| --- | --- | --- | --- | --- |
| 1 | 379 644 | 8214 | 96 | 6 |
| 2 | 621 677 | 5282 | 96 | 6 |
| 3 | 744 749 | 4825 | 95 | 6 |
| 4 | 803 992 | 3723 | 93 | 6 |
| 5 | 580 709 | 7758 | 95 | 6 |
| 6 | 489 588 | 8212 | 95 | 6 |
| 7 | 594 873 | 7244 | 95 | 6 |
| 8 | 639 891 | 5670 | 95 | 6 |
| 9 | 584 711 | 6808 | 95 | 6 |
| 10 | 525 807 | 7029 | 95 | 6 |
| 11 | 831 922 | 3660 | 92 | 6 |
| 12 | 485 181 | 8593 | 96 | 6 |
| 13 | 471 373 | 8815 | 96 | 6 |
| 14 | 792 320 | 3178 | 92 | 6 |
| 15 | 784 016 | 3161 | 92 | 6 |
| 16 | 528 694 | 8417 | 95 | 6 |
| 17 | 423 621 | 9045 | 96 | 6 |
| 18 | 816 536 | 3376 | 93 | 6 |
| 19 | 638 904 | 6031 | 96 | 6 |
| 20 | 770 950 | 4550 | 93 | 6 |
| 21 | 726 080 | 5007 | 95 | 6 |
| 22 | 593 927 | 6227 | 95 | 6 |
| 23 | 617 536 | 5884 | 96 | 6 |
| 24 | 615 115 | 5630 | 95 | 6 |
| 25 | 838 462 | 3504 | 94 | 6 |

## 4 Results

### 4.1 Pair-wise shared SHM are enriched in B cell clones

Clonally related cells will share SHMs that were accumulated by common ancestors over the course of clonal expansion. However, cells from distinct clones are also expected to share mutations at some positions, such as SHM hot-spots (Gadala-Maria *et al.*, 2019). Therefore, we sought to evaluate the degree to which pair-wise shared mutations were enriched in B cell clones. For each simulated dataset, the pair-wise shared SHM matrix *H* was generated for each B cell clone by comparing the IGHV and IGHJ regions of each pair of sequences with the relevant germline sequence. Then, the average of the upper triangular elements was calculated (note that *H* is a symmetric matrix). We found that pair-wise shared SHMs could be identified in ∼ 95% of non-singleton B cell clones (i.e., clones with more than one member) across all simulated datasets. The non-singleton clones without shared mutations tended to be small (with *<* 5 members), so the chance of observing pair-wise shared mutations is lower (Supplementary Figures 1-25C).

We next sought to test whether this high rate of pair-wise SHM sharing was specific to clonally-related sequences. We generated a set of artificial clones (negative controls) by randomly sampling sequences across known clones. Specifically, for each clone from the 100 largest VJ(*ℓ*)-groups (covering ∼ 30% of the total reads), we generated a set of 1000 negative controls with the same size as the given clone. We note that since sampling was performed within each VJ(*ℓ*)-group, the negative controls were generated from sequences with the same junction length, IGHV, and IGHJ genes as the given clone, thus resulting in a conservative control experiment. Then, for each clone and corresponding set of negative controls, the pair-wise shared SHM matrix *H* was generated by comparing the IGHV and IGHJ regions of each pair of sequences with the relevant germline sequence. We performed this analysis for all simulated datasets and calculated the average of the upper triangular elements of *H*. We found that the true clones exhibited significantly (*p <* 0.001) higher pair-wise shared SHM rates (on average ∼ 16 *±* 6 mutations per clone) compared with the set of negative controls (on average ∼ 5 *±* 1 mutations per clone), with a percentage difference of ∼ 105% on average across all simulated datasets (Supplementary Figures 1-25H). Thus, pair-wise shared SHM are enriched in BCR clones. These results support the idea that the pair-wise shared SHM frequency can be leveraged as a biometric (fingerprint) in the clonal relationship inference process.

### 4.2 Focusing on CDR3 improves performance of the recombination-based model

The original recombination-based model for identification of B cell clones used by **SCOPer** measures distance using the junction region of the BCR (Nouri and Kleinstein, 2018b). The junction includes the CDR3 along with the two flanking amino acids (one 5′ that is encoded by IGHV, and one 3′ that is encoded by IGHJ) (Lefranc, 2014). As the two flanking positions are highly conserved, we sought to determine whether they were necessary to include in the distance measure. Indeed, we hypothesized that including these positions could even lead to decreased performance, as they are likely to be identical across independent clones and will have increasing influence on the distance for clones with shorter junction lengths.

To test this hypothesis, we compare the performance of the recombination-based model using either the junction-targeted (termed as `ham-junc`) or CDR3-targeted (termed as `ham-cdr3`) approaches. Using simulated data, performance was quantified using the measures of sensitivity, specificity, and precision. The sensitivity (true positive rate) of each model is defined as the fraction of all sequence pairs from the same clone that were correctly inferred by the model, while specificity (true negative rate) is defined as the fraction of pairs of unrelated sequences that were successfully inferred by the model to be in different clones. Finally, the precision (positive predictive value) of each model is defined by measuring how often inferred clonal relative sequence pairs are truly clonally related. We found that both approaches inferred the clonal relationships with high sensitivity, specificity, and precision with values of *>* 94.0% on average across all simulated datasets. However, each of the measures of accuracy were significantly (*p <* 0.001) improved when distance was based on the CDR3 region, rather than the junction region (Figure 5). Thus, the conserved positions flanking the junction should not be used to define the distance between sequences.

**Fig. 5.**
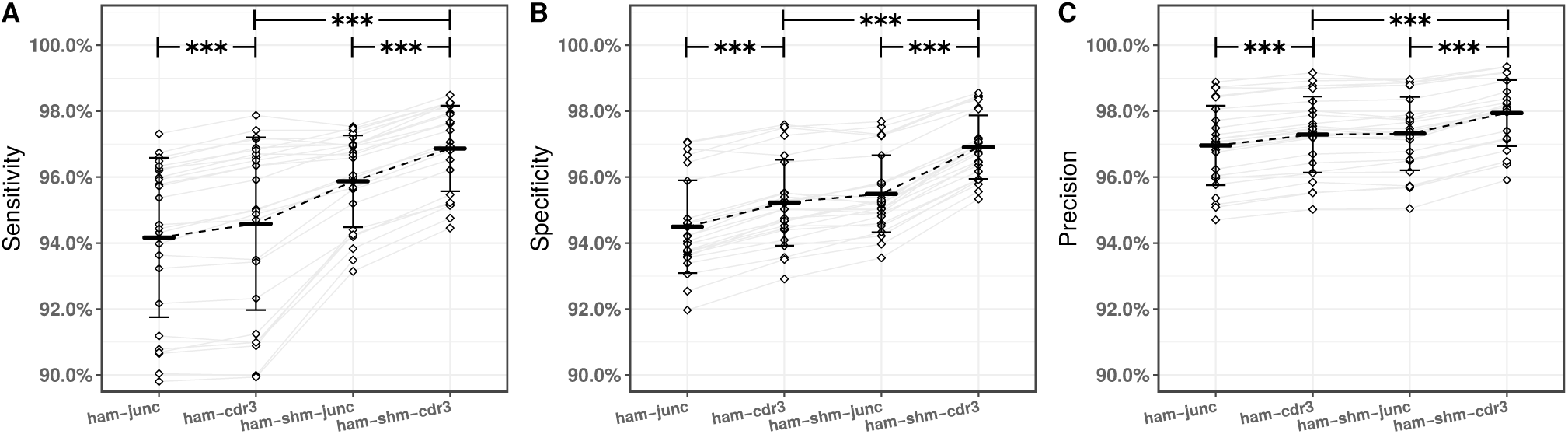
Integrating information from CDR3 similarity (recombination-based distance) and shared mutations in the V and J segments (SHM-based distance) improves clonal relationship inference. The spectral clustering-based framework was applied to identify clonally-related sequences in 25 simulated datasets (diamonds) generated via **AbSim** R package (Yermanos et al., 2017) (Table 1). Performance was assessed by calculating (A) sensitivity, (B) specificity, and (C) precision via applying the recombination-based model on the junction (`ham-junc`) and CDR3 (`ham-cdr3`) regions, as well as the integrated model on the junction (`ham-shm-junc`) and CDR3 (`ham-shm-cdr3`) regions. Mean performance is indicated by the solid bars, while the error bars define one standard deviation. For the comparisons of interest the asterisks (*\*\*\**) indicate *p <* 0.001 by paired t-test.

### 4.3 Shared mutations should be integrated with CDR3 distance to identify clones

We next asked whether incorporating shared SHMs of V and J segments into the procedure leads to even better performance. We thus characterized the performance of integrated model using CDR3-targeted (termed as `ham-shm-cdr3`) approach. Including shared SHM with the integrated model improved measures of sensitivity, specificity, and precision to ≳ 97% on average across all simulated datasets. For the sake of completeness, we also characterized the performance of integrated model using junction-targeted (termed as ham-shm-junc) approach. Consistent with our analysis of the recombination-based model, we found that using the junction rather than the CDR3 region led to a significant (*p <* 0.001) decrease in performance (Figure 5).

These results indicate that the best performance within the spectral clustering-based framework is achieved when the integrated model was accompanied with a CDR3-targeted approach. Overall, when the original **SCOPer** model (`ham-junc`) is compared to the new integrated model (`ham-shm-cdr3`), a ∼ 3% improvement in the sensitivity, ∼ 2.5% improvement in the specificity, and ∼ 1% improvement in the precision was achieved on average across all simulated datasets (Figure 5).

To better understand how the integrated model improves the performance of clonal relationship inference, we examined its operation in detail using one of the identified VJ(*ℓ*)-groups with 42 unique sequences. As these are simulated data, we know that these sequences are comprised of two clones, one consisting of 41 sequences, and the other of only one sequence. Comparing the clonal relationships using the CDR3-targeted recombination-based model (`ham-cdr3`) and the CDR3-targeted integrated model (`ham-shm-cdr3`), we find that both models inferred two clones. However, `ham-cdr3` model failed to accurately infer the clonal relationships of one of the sequences, which resulted in one false positive and multiple false negatives (Figure 6A). On the other hand, when the SHM among sequences was expressed using the pair-wise SHMs (on average ∼ 23 *±* 8 mutations were counted per pair, from which ∼ 6 *±* 5 mutations were shared), the clonally-related sequences were pulled toward each other whereas the singleton (with at least 2-fold fewer shared pair-wise mutation than other clone members) remained separated, thereby the performance of the local scaling procedure was improved (Figure 6B). Hence, the `ham-shm-cdr3` model resulted in no false relationships in this particular case (Figure 6C).

**Fig. 6.**
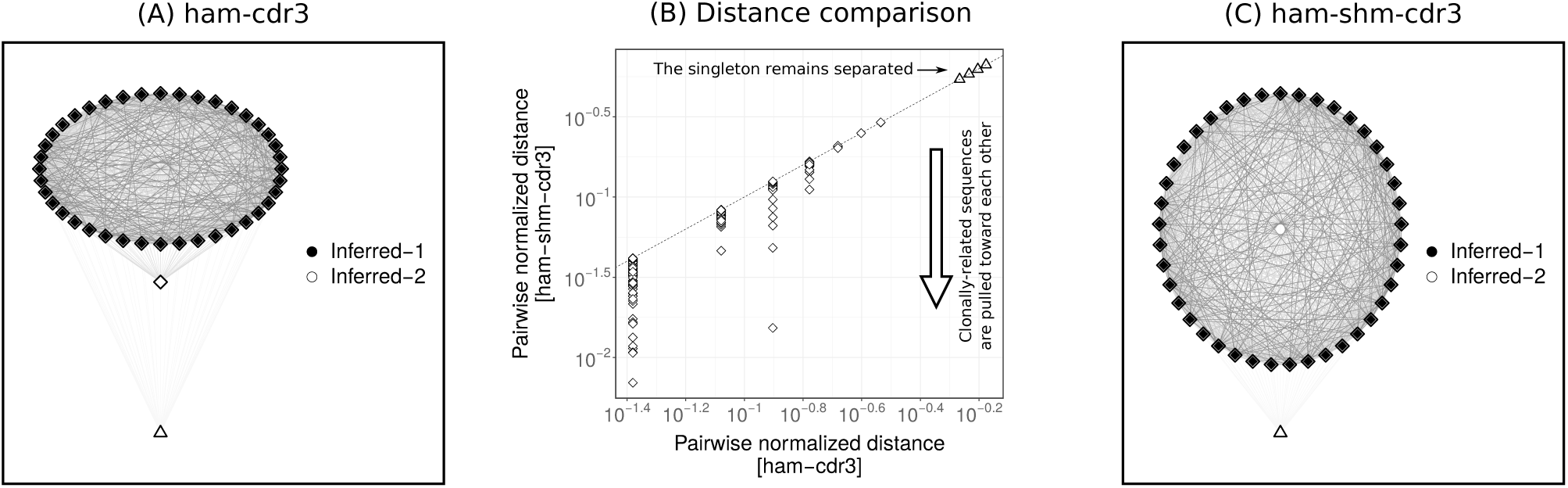
The integrated model improves clonal inference by pulling clonally-related sequences toward each other. The spectral clustering-based model was applied to infer the clonal relationships among 42 sequences from a given VJ(*ℓ*)-group. These sequences belong to two clones, one consisting of 41 sequences (diamonds), and the other only one sequence (triangle). Clonal relationships were inferred (indicated by filled colors) via the CDR3-targeted recombination-based model (`ham-cdr3`) leading to two clones (Inferred-1 and Inferred-2) (A), and CDR3-targeted integrated model (`ham-shm-cdr3`) leading to two clones (Inferred-1 and Inferred-2) (C). For visualization, the sequences were embedded in 2D space using the `qgraph` function from the **qgraph** R package, where the thickness of each edge indicates the inverse of the pair-wise `ham-cdr3` (A) and `ham-shm-cdr3` distances (C). Pair-wise distances were normalized by the CDR3 length and compared in log scale (B).

### 4.4 The integrated model performs with high confidence on experimental data

Along with simulated data, we also evaluated the performance of the CDR3-targeted integrated model (`ham-shm-cdr3`) by estimating specificity using experimental BCR sequencing data from 58 individuals with acute dengue infection (Parameswaran *et al.*, 2013). By definition, clones cannot span different individuals. To estimate specificity, we combine data from multiple individuals, use the `ham-shm-cdr3` model to identify clonal relationships, and then count the frequency of clones that are (incorrectly) inferred to be shared across individuals (Gupta *et al.*, 2017). We use the procedure proposed in (Nouri and Kleinstein, 2018a). First, one of the individuals (the dataset with largest number of unique sequences) was chosen as the “base”. Next, a single sequence was chosen randomly from each of the remaining individuals and added to the sequencing data from the base individual. Specificity was then defined by how often the sequences from non-base individuals were correctly determined to be singletons. Any grouping of these sequences into larger clones must be a false positive. This procedure was then repeated for 100 cycles. The results indicated that the `ham-shm-cdr3` model has a high specificity with a value of ∼ 96.0% on average across all cycles. Thus, combining shared SHMs in the V and J segments of the BCR can be leveraged along with the CDR3 sequence to identify clonally related sequences with high specificity in experimental data.

### 4.5 The SCOPer algorithm is efficiently parallelized

Computational efficiency is an important property considering the recent growth in the size of typical BCR repertoires (Soto *et al.*, 2019; Briney *et al.*, 2019). Using the recombination-based model we found that clonal partitioning ∼ 640 *±* 95k simulated sequences (the average repertoire size used in this study) took ∼ 30 *±* 5 min, but when the integrated model was involved the partitioning took ∼ 160 *±* 15 min. This assessment was performed on a Linux computer with a 2.20 GHz Intel processor and 32 GB RAM. There are two main factors that drive this increased computational cost. In our current implementation, clonal inference is performed on the set of unique sequences (i.e., sequences with distinct nucleotide sequences). When using a recombination-based model that considers only the junction or CDR3, the chance of having identical sequences in each VJ(*ℓ*)-group is high (on average across all simulated datasets ∼ 60% of CDR3s are unique per each VJ(*ℓ*)-group). This decreases the computational cost of the algorithm. In contrast, when using the integrated model, the V and J segments are also relevant, allowing fewer sequences to be combined into identical groups (i.e., leading to more unique sequences). The computational cost increases with this increasing number of sequences *n*. Specifically, the eigen-decomposition algorithm, which scales by 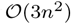 (we note that the targeted matrix, to be spectrally decomposed, is symmetric which improves the computational cost significantly). Furthermore, the pair-wise SHM analysis brings additional computational complexity. For instance, the computational complexity of generating the pair-wise shared SHM matrix *H* algorithm is 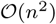. This run time will be summed up by the pair-wise recombination-based matrix *X* with the same computational complexity. However, the **SCOPer** distributed implementation facilitates the clonal inference process by parallelizing the computation and greatly reducing the running time. In our current implementation, the parallelization is achieved by distributing the clonal inference process from each VJ(*ℓ*)-group of sequences across processing cores dynamically. The parallelization is possible on cores from a single workstation or on high-performance computing (HPC) cluster facilities. For instance, using only five cores in parallel decreased the running time to ∼ 44 *±* 11 min, a ∼ 4-fold improvement, for partitioning ∼ 640 *±* 95k sequences. Our benchmarks across all simulated data sets demonstrate good scalability resulting in a speedup, defined as the time it takes the integrated algorithm to execute with one processor divided by the time it takes to execute in parallel, that is approximately linear to the number of cores (*<* 10) utilized (Figure 7).

**Fig. 7.**
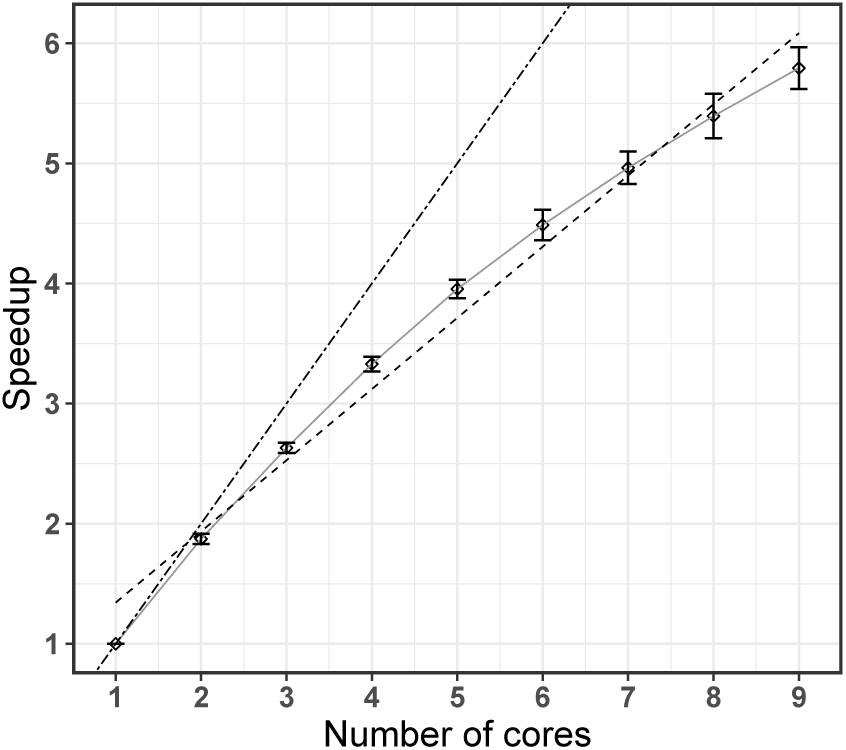
The **SCOPer** algorithm can be run efficiently on multiple cores. The speedup, defined as the time it takes the algorithm to execute with one processor divided by the time it takes to execute in parallel, was calculated for the integrated model for different numbers of processing cores. In each case, speedup was calculated as the average across 25 simulated data sets (with error bars showing the standard deviation). Evaluation was carried out on a Linux computer with a 2.20 GHz Intel processor and 32 GB RAM. The linear fit is shown by a dashed line, while the ideal speedup is shown by the dot-dash line.

## 5 Conclusion

B cell clonal diversity is introduced through two main mechanisms. The first occurs during maturation in the bone marrow by stochastic joining of germline-encoded V, D, and J heavy chain genes (or V and J light chain genes) combined with the action of exonucleases and terminal deoxynucleotidyl transferase, which add diversity at the recombination boundaries. This diversity acts as a fingerprint that can be used to separate distinct clones based on the distance between their junction (or CDR3) nucleotide sequence (inter-clonal diversity). Subsequently, upon encountering cognate antigen, B cells can enter a germinal center and undergo further diversification through SHM and affinity maturation. The accumulation of SHMs has the effect of spreading out the sequences of B cell clonal variants around their initial points of creation (intra-clonal diversity). A significant challenge in the clonal relationships inference problem is to define meaningful metrics which can leverage inter-clonal diversity to recognize sequences that are part of independent clones (specificity), while also modeling intra-clonal diversity to recognize the variants that are clonally-related (sensitivity).

We developed an unsupervised learning algorithm based on spectral clustering that provides a framework for the inference of B cell clonal relationships. This model combines CDR3 similarity with shared SHM profiles in the V and J segments to capture both inter- and intra-clonal diversification. We showed that the inclusion of pair-wise shared SHM patterns improves the models ability to identify clonally related sequences. Overall, the model determines B cell clones by: (1) common IGHV- and IGHJ-gene calls and identical CDR3 length, (2) identical or similar CDR3 nucleotide sequences, and (3) shared somatic hypermutation patterns in the V and J segments.

In the absence of gold standard experimental data with known clonal relationship between sequences, the validation was performed using B cell simulations which offer a mechanism to generate data where the underlying clonal groups are known. However, using experimental data we also reported a measure of specificity based on the frequency of clones that are predicted to be shared across individuals.

The influence of SHM hot- and cold-spot biases in the clonal inference process have been incorporated using an SHM targeting model. The analysis described here uses the S5F targeting model for SHM that was previously constructed (Yaari *et al.*, 2013). However, while hot- and cold-spot biases are generally conserved across individuals, these intrinsic biases can be altered by age (Hoehn *et al.*, 2019), and may also differ across species (Cui *et al.*, 2016). Clonal identification could be improved by using a data-specific targeting model that can be built using toolkits available in the Immcantation framework (www.immcantation.org). The S5F model seeks to avoid the biases introduced by selection, and to capture only the intrinsic biases introduced by the activation-induced cytidine deaminase (AID) binding preferences and error-prone DNA repair in a 5-mer micro-sequence context (Yaari *et al.*, 2013). Future improvements to the SHM targeting model, such as including effects beyond motif-specificity (MacCarthy *et al.*, 2009), may also improve clonal relationship inference. However, these must be rigorously tested.

While the model presented here was developed and tested for sequencing data from the H chain only, cutting-edge technologies, including single-cell sequencing, provide paired IGH- and IGL-chain data (DeKosky *et al.*, 2015; Macosko *et al.*, 2015; Briggs *et al.*, 2017). These paired data can be incorporated into the proposed model by extending the criteria for the initial grouping of sequences (i.e., VJ(*ℓ*)-groups) to include the same IGHV-gene, IGHJ-gene, IGH-CDR3 length, IGLV-gene, IGLJ-gene, and IGL-CDR3 length. BCR clonal inference can then be carried out as before on the H chain of these more refined groups. The low diversity of the IGL-chain junction region makes it unlikely that including this region in the clustering will provide a significant performance improvement (Zhou and Kleinstein, 2019).

The definition of clone used in this work is based on the assumption that SHM introduces only point substitutions into the BCR sequence. However, it has been shown that insertions and deletions (indels) can also be introduced at a low frequency (*<* 2-3%) (Smith *et al.*, 1996; Ohlin and Borrebaeck, 1998; Wilson *et al.*, 1998; de Wildt *et al.*, 1999; Briney *et al.*, 2012; Hwang *et al.*, 2017). Distance functions that allow for sequences of different lengths could be used to identify clonally related sequences that differ by indels (leading, for example, to sequences with different CDR3 lengths). However, these must be rigorously tested.

The models described in this study have been implemented in the **SCOPer** (Spectral Clustering for clOne Partitioning) R package, which provides a computational framework to explore multiple approaches to infer clonal relationships in AIRR-seq data. This implementation of **SCOPer** is freely available as part of the Immcantation framework (www.immcantation.org) under the CC BY-SA 4.0 license. The input and output formats of **SCOPer** conform to the Change-O (Gupta *et al.*, 2015) and AIRR (Vander Heiden *et al.*, 2018) file standard, and thus the method can be used seamlessly as part of the Immcantation tool suite, including methods for B cell clonal lineage reconstruction, lineage topology analysis, clonal diversity analysis, and other advanced repertoire analyses linked to the clonal landscape.

## 6 Data availability

The simulated data are accessible at http://clip.med.yale.edu/papers/Nouri2019A.

## Acknowledgements

This work was supported in part by the National Institutes of Health (NIH) under award number R01AI104739 and a National Library of Medicine (NLM) Fellowship. We thank the Yale Center for Research Computing for guidance and use of the research computing infrastructure. We wish to acknowledge Alina Aleksandrova for a careful reading of the manuscript. The authors also thank Susanna Marquez for useful comments related to the development of the code.

## Author contributions

N.N. and S.H.K. conceived and designed the project. N.N. implemented the method. Both authors wrote and edited the manuscript.

