## Supplementary Figures for "Somatic hypermutation analysis for improved identification of B cell clonal families from next-generation sequencing data"

**Supplementary figure 1:** The global metrics of the BCR simulated repertoires, including: (1) junction length distribution, (2) distance-to-nearest distribution, (3) clonal relative abundance distribution, (4) clone size distribution, (5) mutation frequency distribution, (6) number of clones per VJℓ-group, (7) average pair-wise SHM for clone, and (8) negative-control test (comparing pair-wise SHM sharing rate among real clones and a set of artificial clones generated by randomly sampling sequences across known clones).

**Simulation-1**

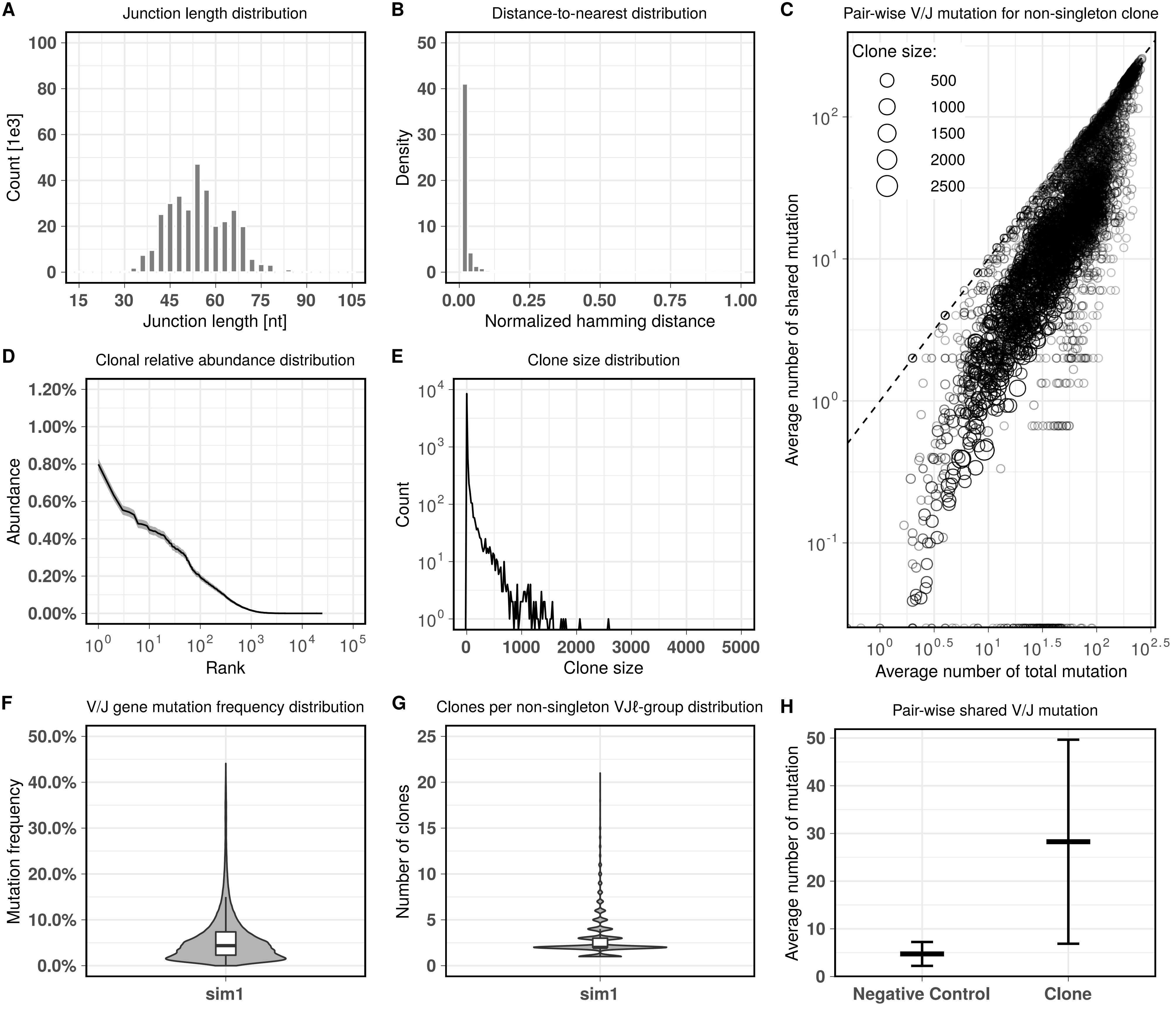

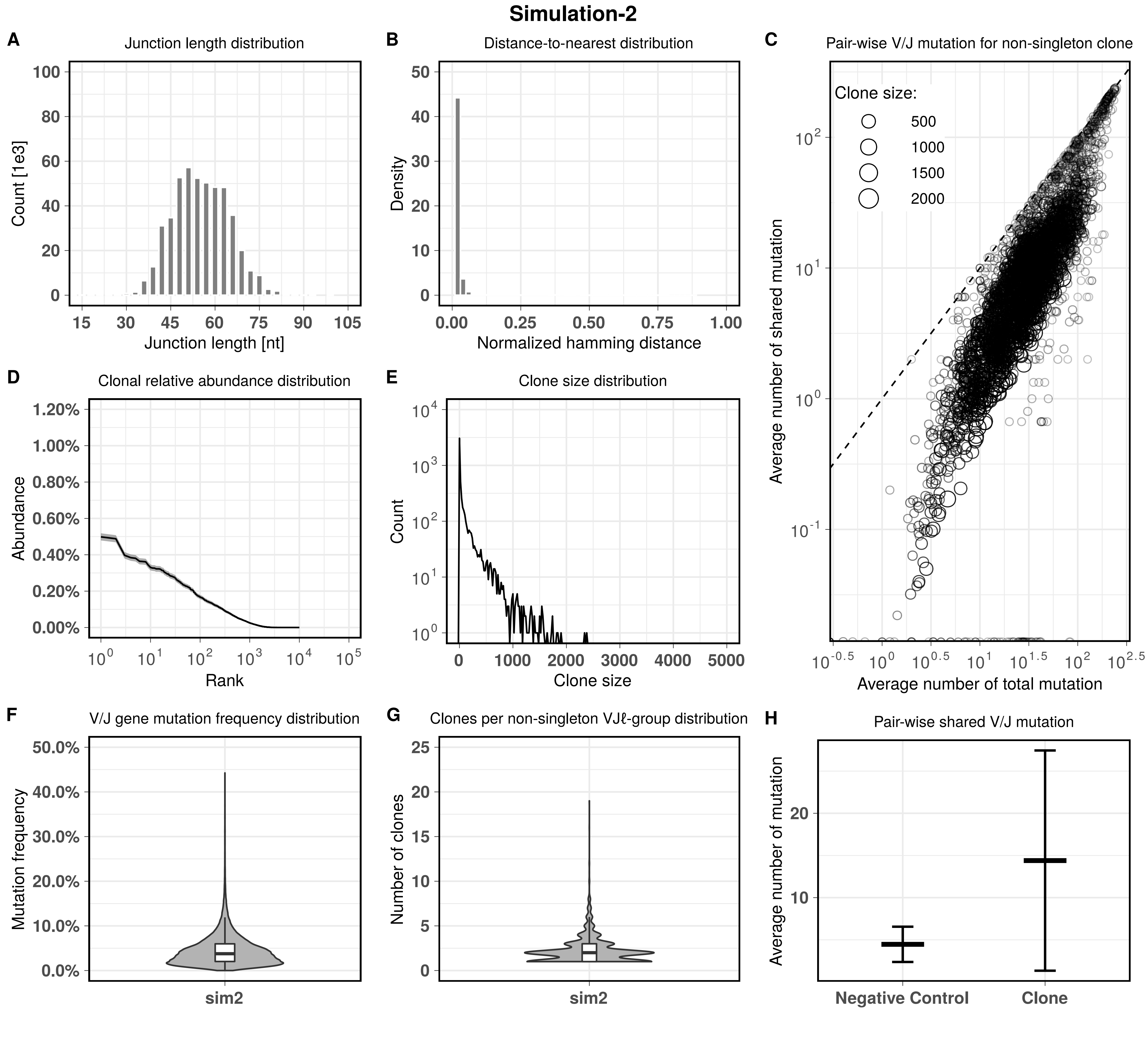

Simulation-2
A
Junction length distribution
Count [1e3]
Junction length [nt]
B
Distance-to-nearest distribution
Density
Normalized hamming distance
C
Pair-wise V/J mutation for non-singleton clone
Clone size:
500
1000
1500
2000
Average number of shared mutation
Average number of total mutation
D
Clonal relative abundance distribution
Abundance
Rank
E
Clone size distribution
Count
Clone size
F
V/J gene mutation frequency distribution
Mutation frequency
sim2
G
Clones per non-singleton VJℓ-group distribution
Number of clones
sim2
H
Pair-wise shared V/J mutation
Average number of mutation
Negative Control
Clone

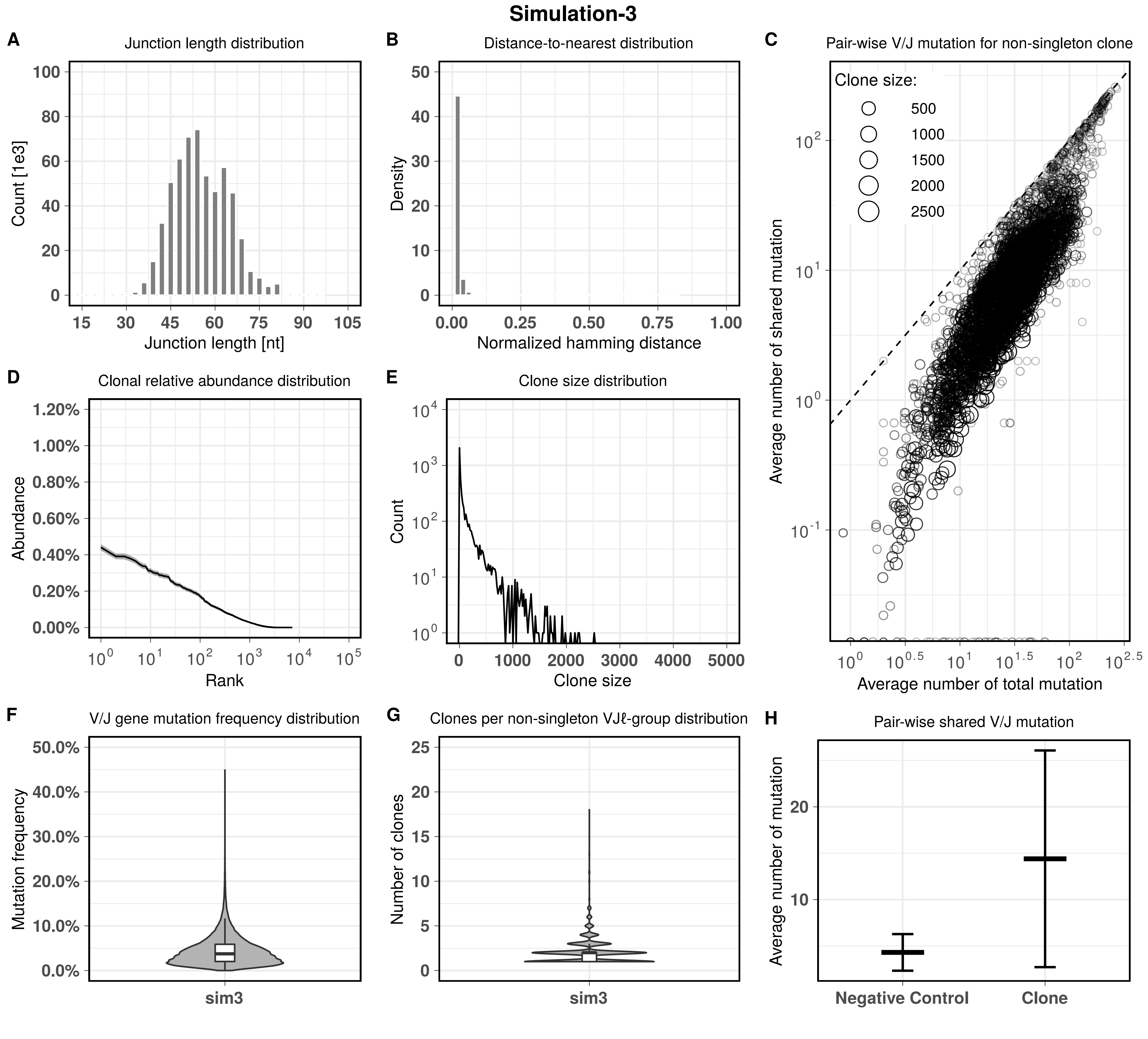

Simulation-3
A
Junction length distribution
Count [1e3]
Junction length [nt]
B
Distance-to-nearest distribution
Density
Normalized hamming distance
C
Pair-wise V/J mutation for non-singleton clone
Clone size:
500
1000
1500
2000
2500
Average number of shared mutation
Average number of total mutation
D
Clonal relative abundance distribution
Abundance
Rank
E
Clone size distribution
Count
Clone size
F
V/J gene mutation frequency distribution
Mutation frequency
sim3
G
Clones per non-singleton VJℓ-group distribution
Number of clones
sim3
H
Pair-wise shared V/J mutation
Average number of mutation
Negative Control
Clone

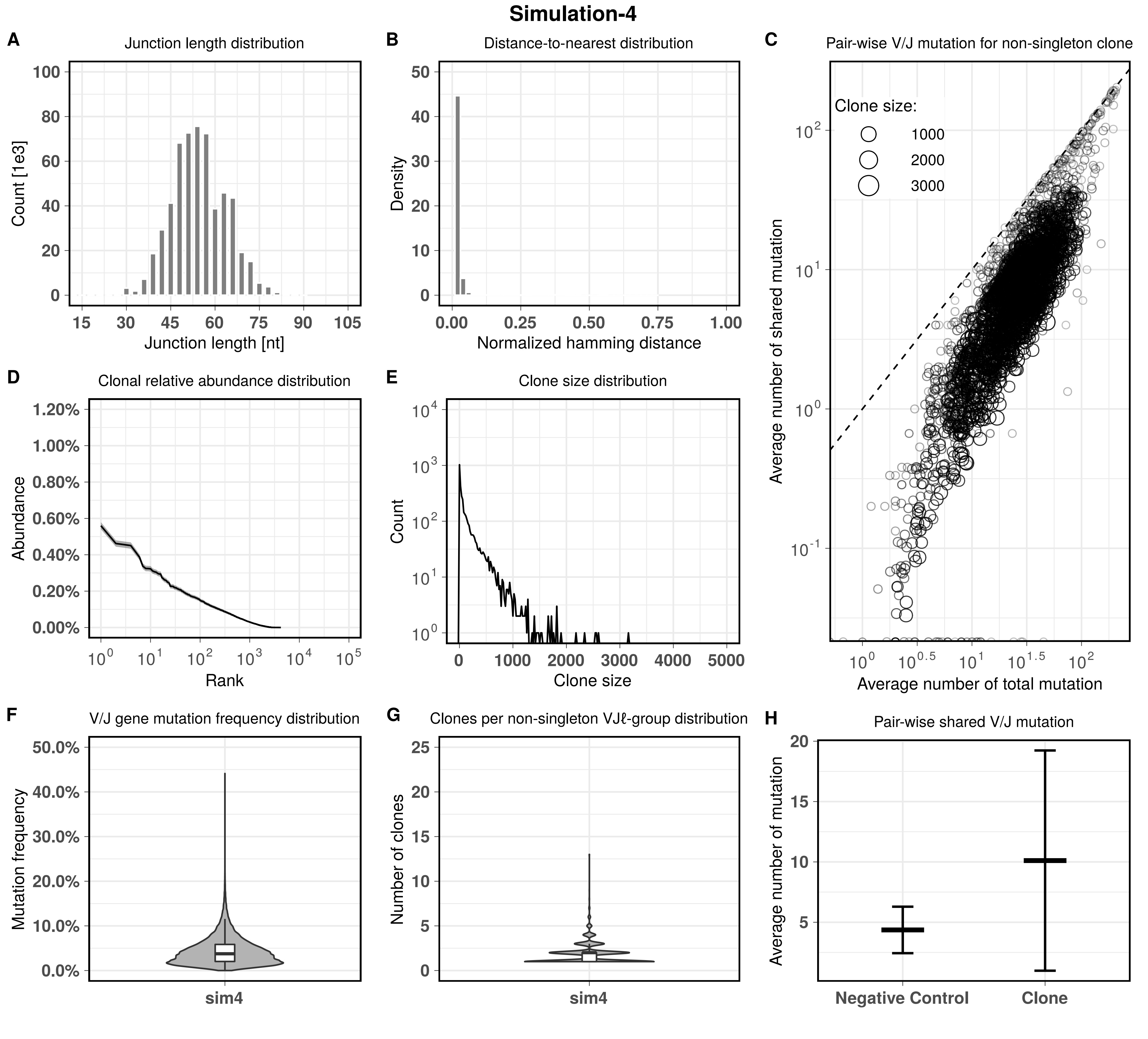
Simulation-4
A
Junction length distribution
Count [1e3]
Junction length [nt]
B
Distance-to-nearest distribution
Density
Normalized hamming distance
C
Pair-wise V/J mutation for non-singleton clone
Clone size:
1000
2000
3000
Average number of shared mutation
Average number of total mutation
D
Clonal relative abundance distribution
Abundance
Rank
E
Clone size distribution
Count
Clone size
F
V/J gene mutation frequency distribution
Mutation frequency
sim4
G
Clones per non-singleton VJℓ-group distribution
Number of clones
sim4
H
Pair-wise shared V/J mutation
Average number of mutation
Negative Control
Clone

#### Simulation-5

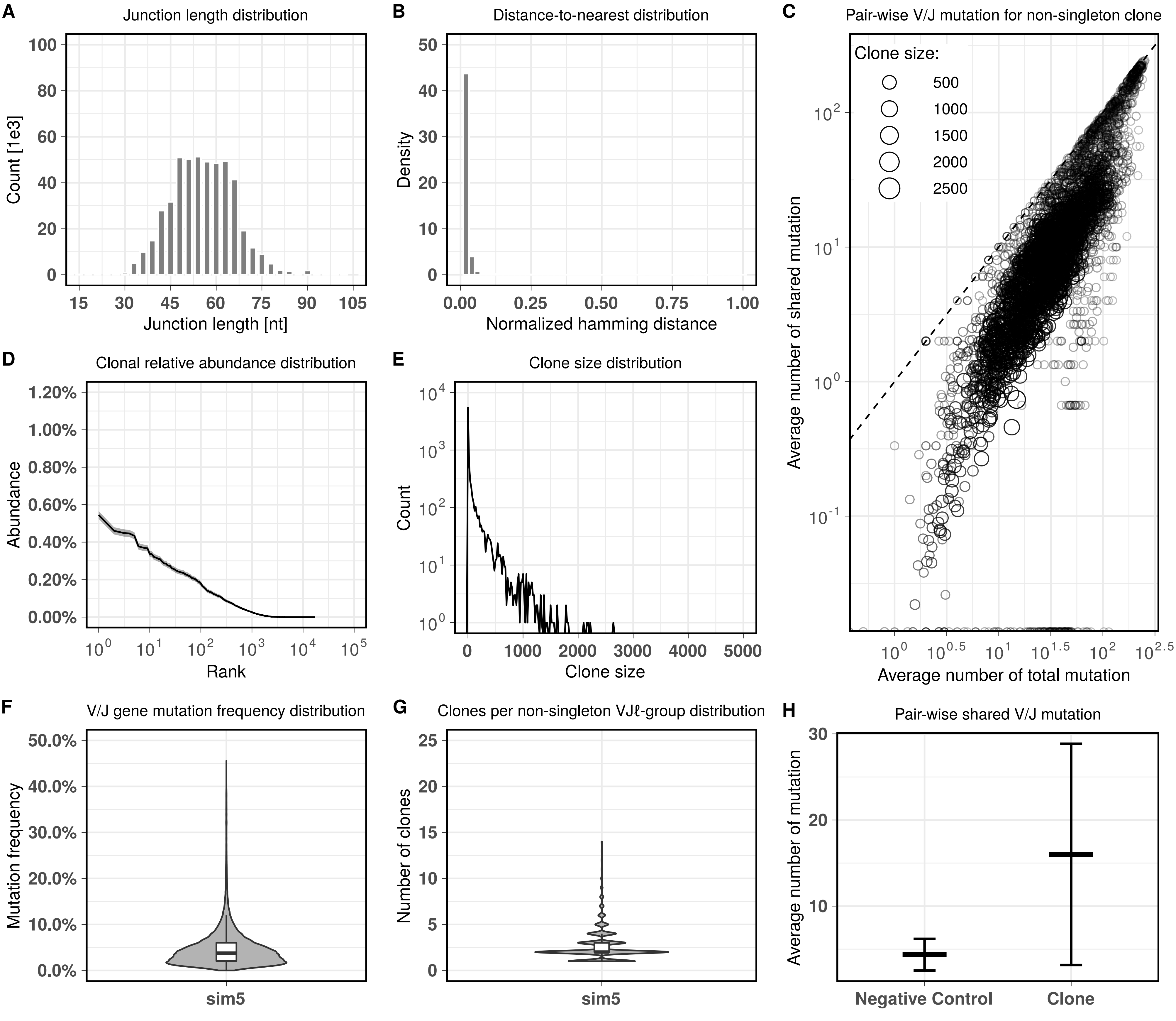

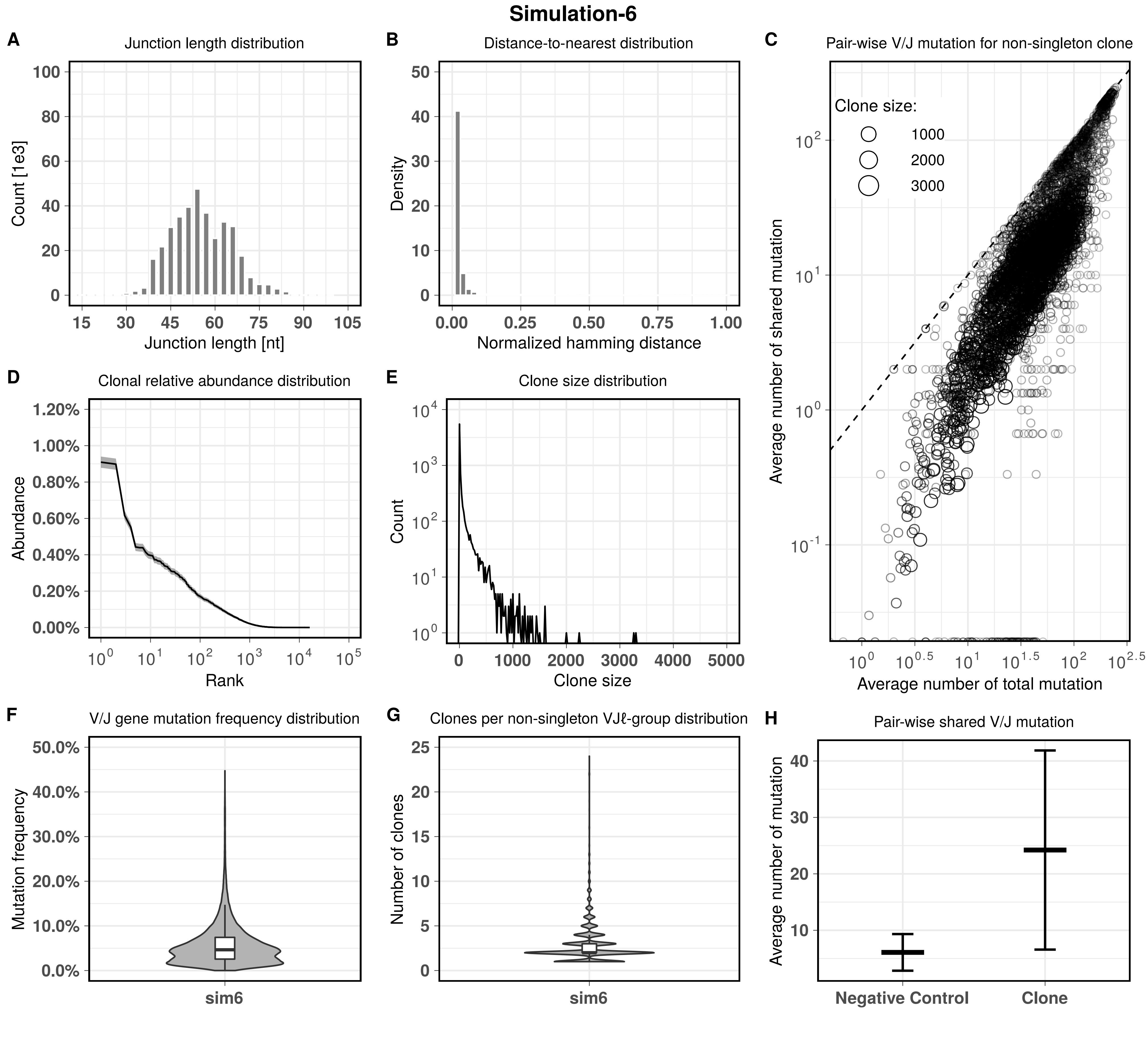
Simulation-6
A
Junction length distribution
Count [1e3]
Junction length [nt]
B
Distance-to-nearest distribution
Density
Normalized hamming distance
C
Pair-wise V/J mutation for non-singleton clone
Clone size:
1000
2000
3000
Average number of shared mutation
Average number of total mutation
D
Clonal relative abundance distribution
Abundance
Rank
E
Clone size distribution
Count
Clone size
F
V/J gene mutation frequency distribution
Mutation frequency
sim6
G
Clones per non-singleton VJℓ-group distribution
Number of clones
sim6
H
Pair-wise shared V/J mutation
Average number of mutation
Negative Control
Clone

**Simulation-7**

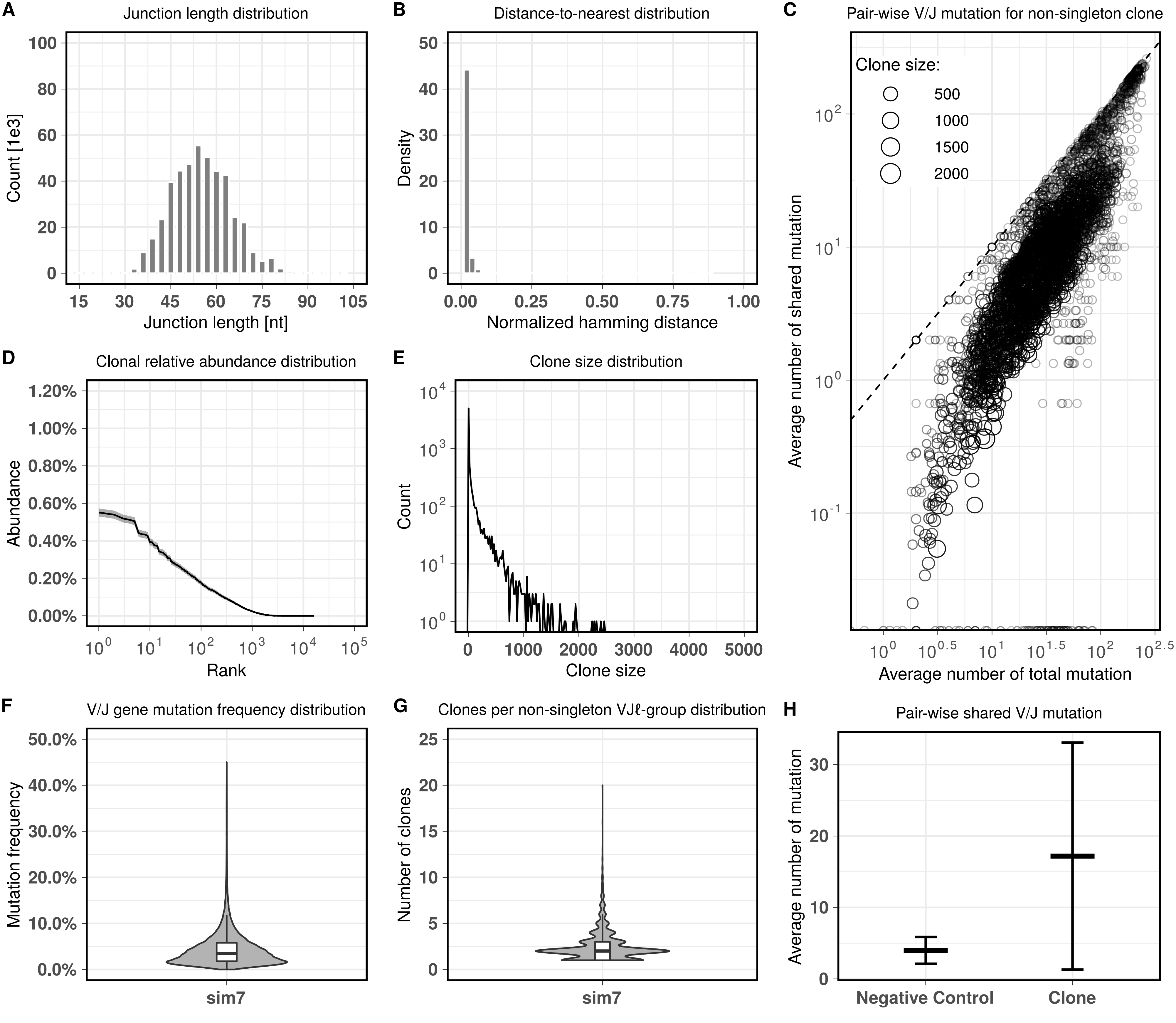

#### Simulation-8

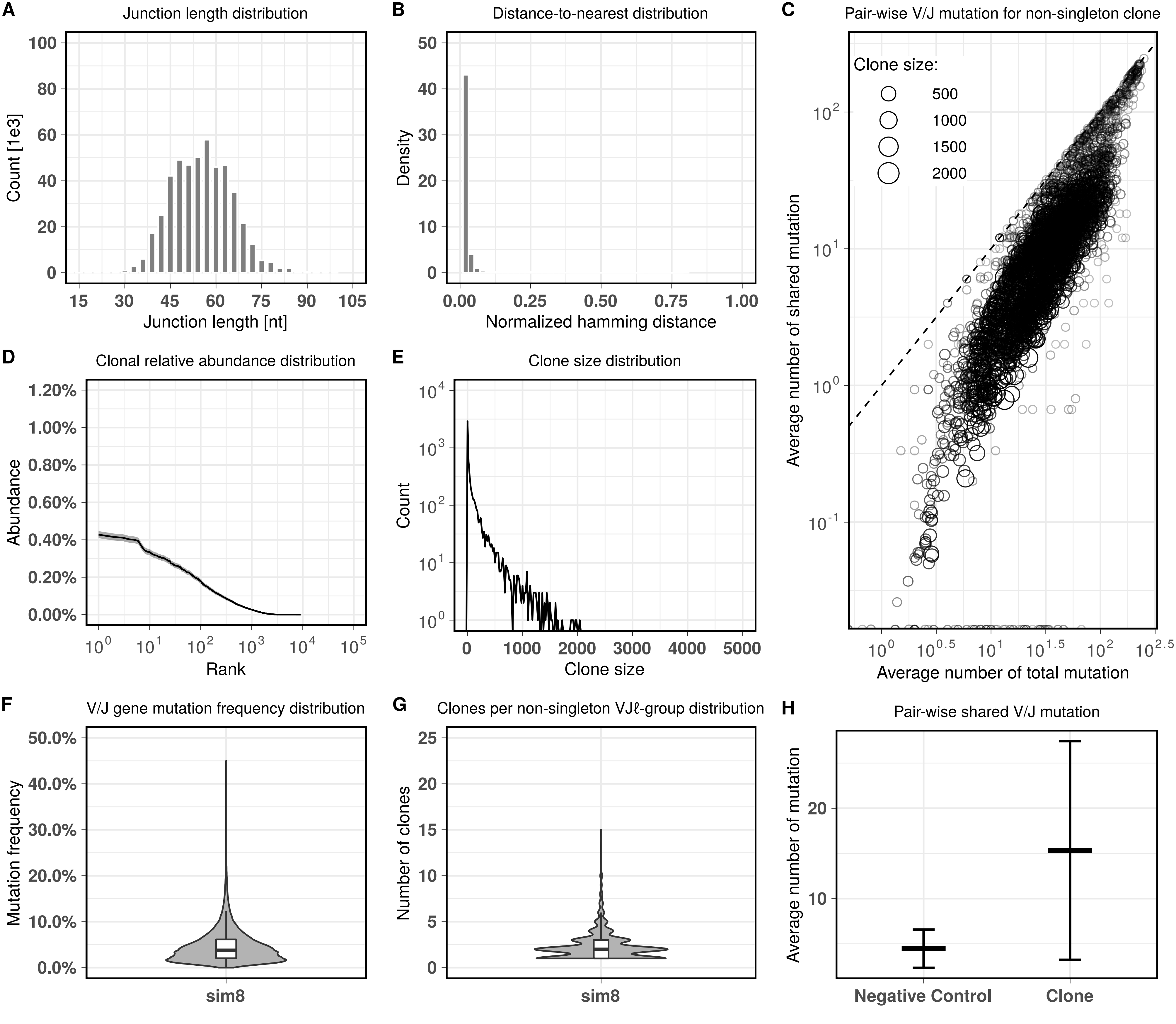

**Simulation-9**

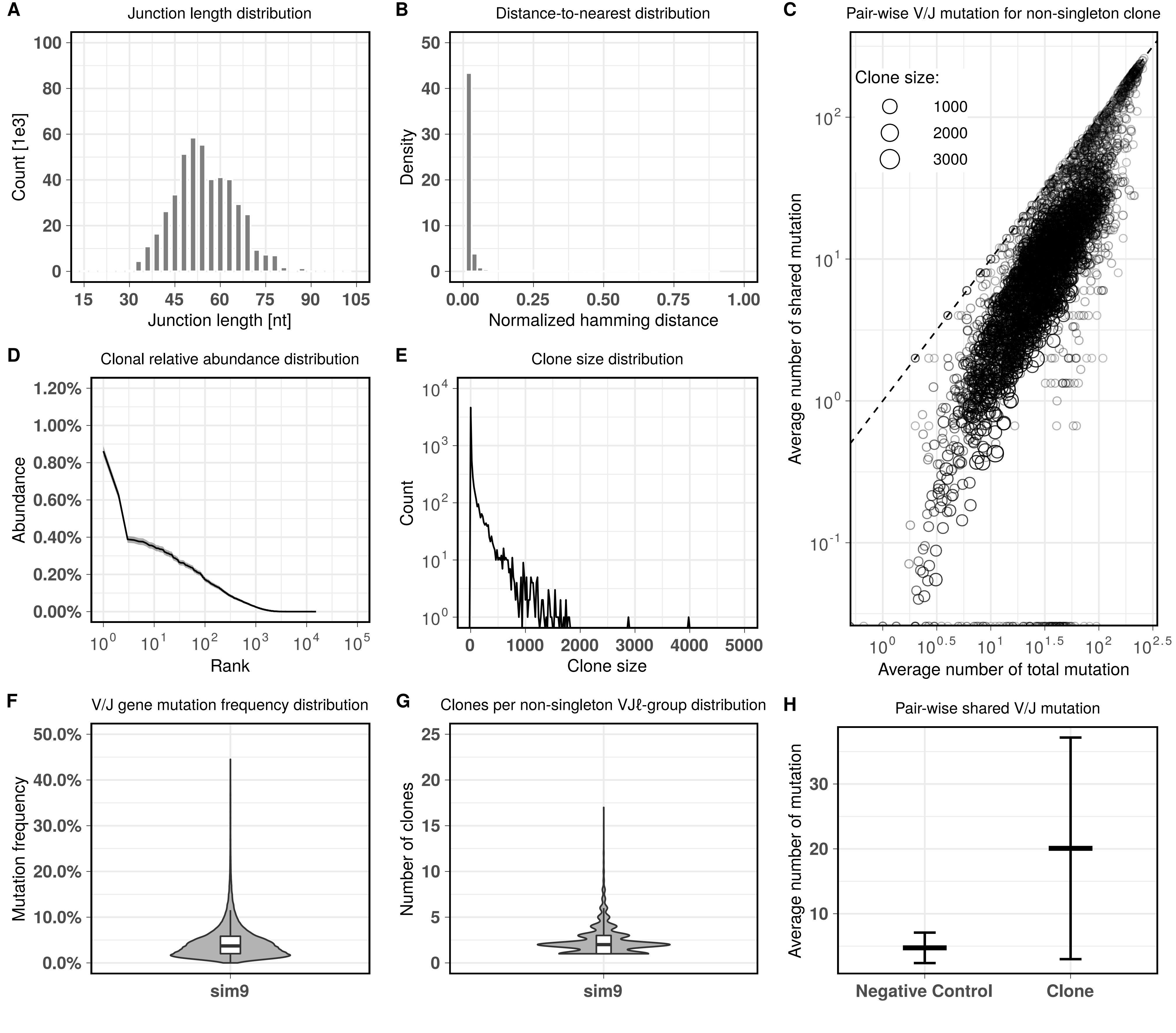

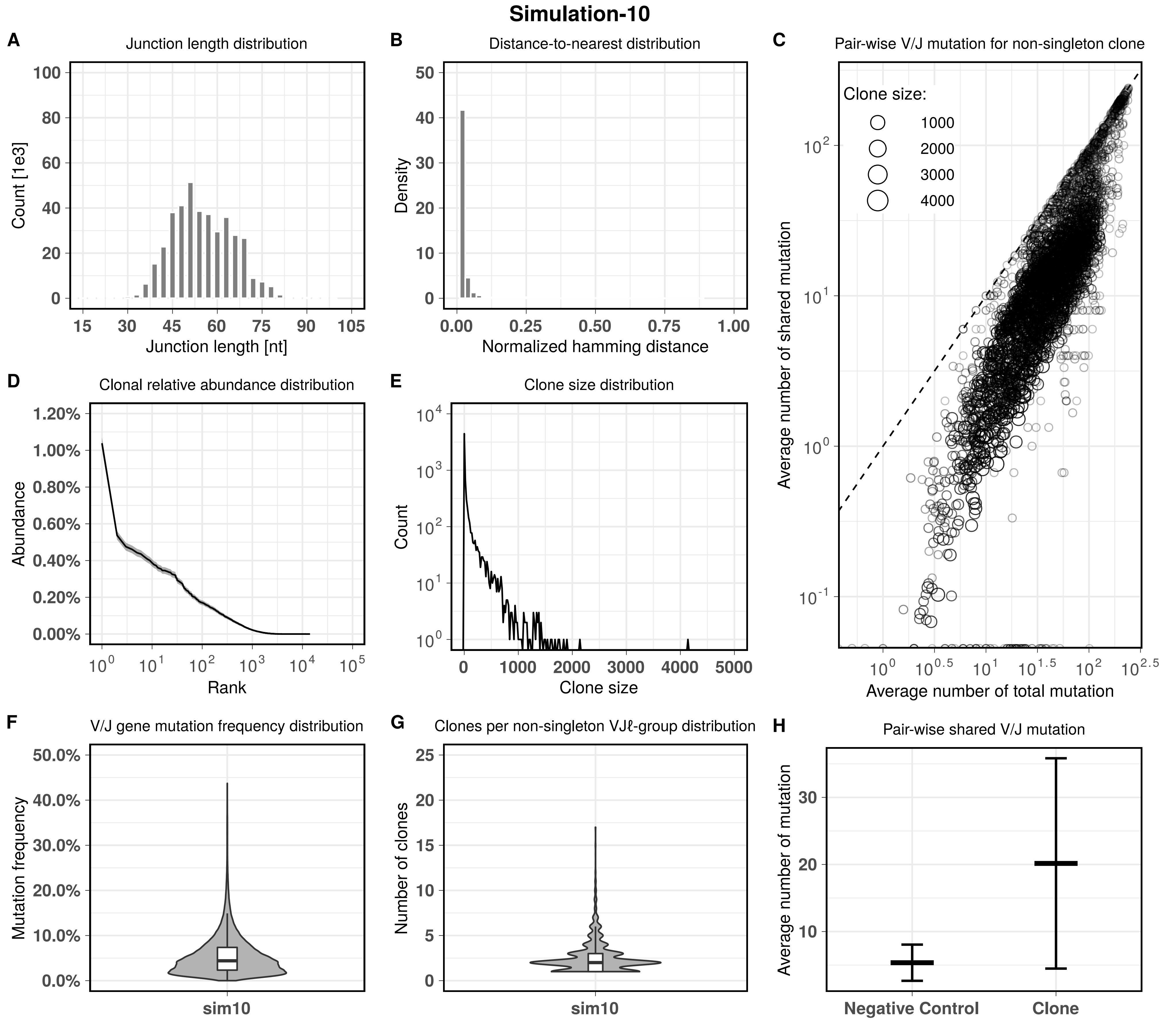

Simulation-10
A
Junction length distribution
Count [1e3]
Junction length [nt]
B
Distance-to-nearest distribution
Density
Normalized hamming distance
C
Pair-wise V/J mutation for non-singleton clone
Clone size:
1000
2000
3000
4000
Average number of shared mutation
Average number of total mutation
D
Clonal relative abundance distribution
Abundance
Rank
E
Clone size distribution
Count
Clone size
F
V/J gene mutation frequency distribution
Mutation frequency
sim10
G
Clones per non-singleton VJℓ-group distribution
Number of clones
sim10
H
Pair-wise shared V/J mutation
Average number of mutation
Negative Control
Clone

### Simulation-11

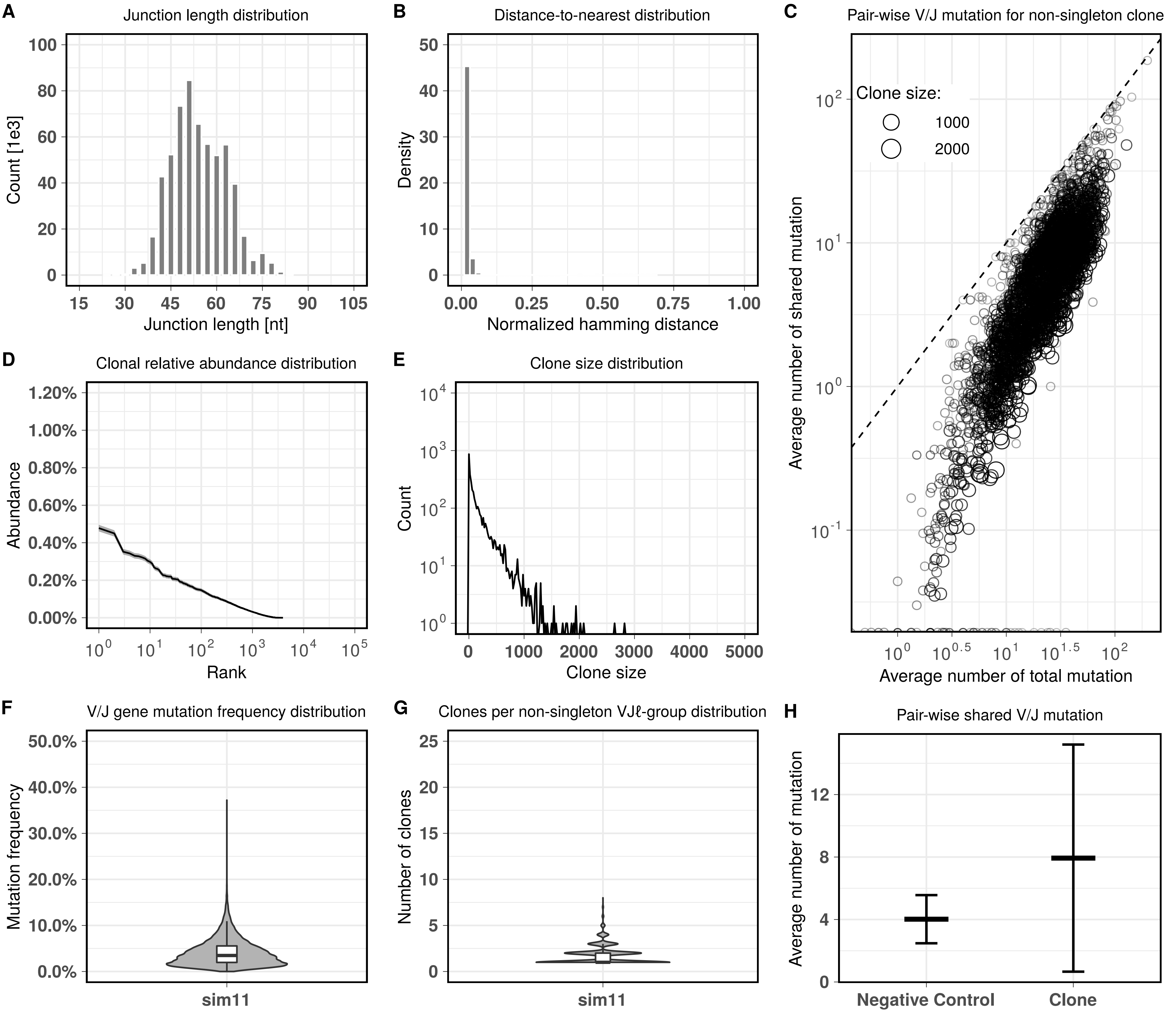

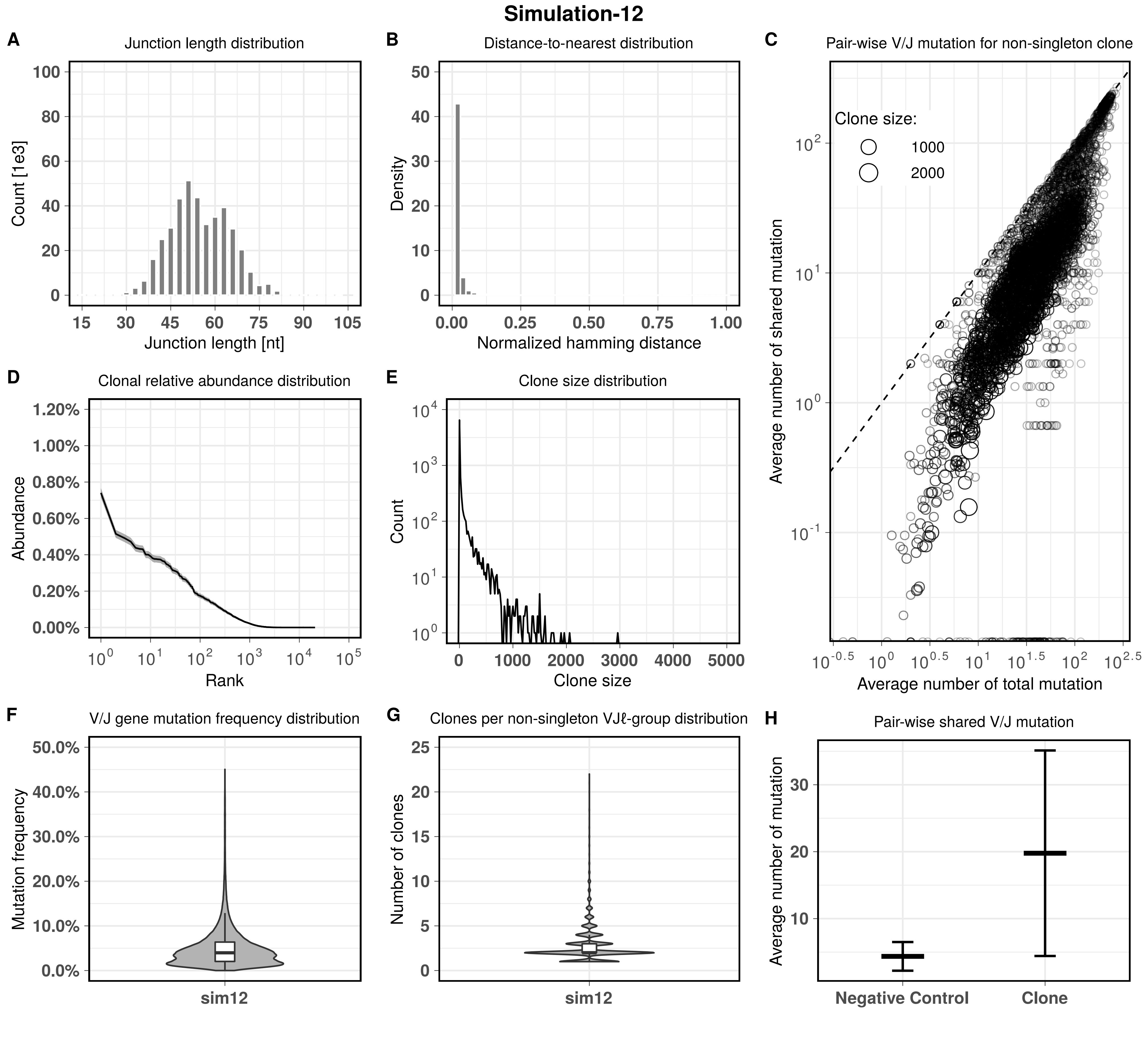
Simulation-12
A
Junction length distribution
Count [1e3]
Junction length [nt]
B
Distance-to-nearest distribution
Density
Normalized hamming distance
C
Pair-wise V/J mutation for non-singleton clone
Clone size:
1000
2000
Average number of shared mutation
Average number of total mutation
D
Clonal relative abundance distribution
Abundance
Rank
E
Clone size distribution
Count
Clone size
F
V/J gene mutation frequency distribution
Mutation frequency
sim12
G
Clones per non-singleton VJℓ-group distribution
Number of clones
sim12
H
Pair-wise shared V/J mutation
Average number of mutation
Negative Control
Clone

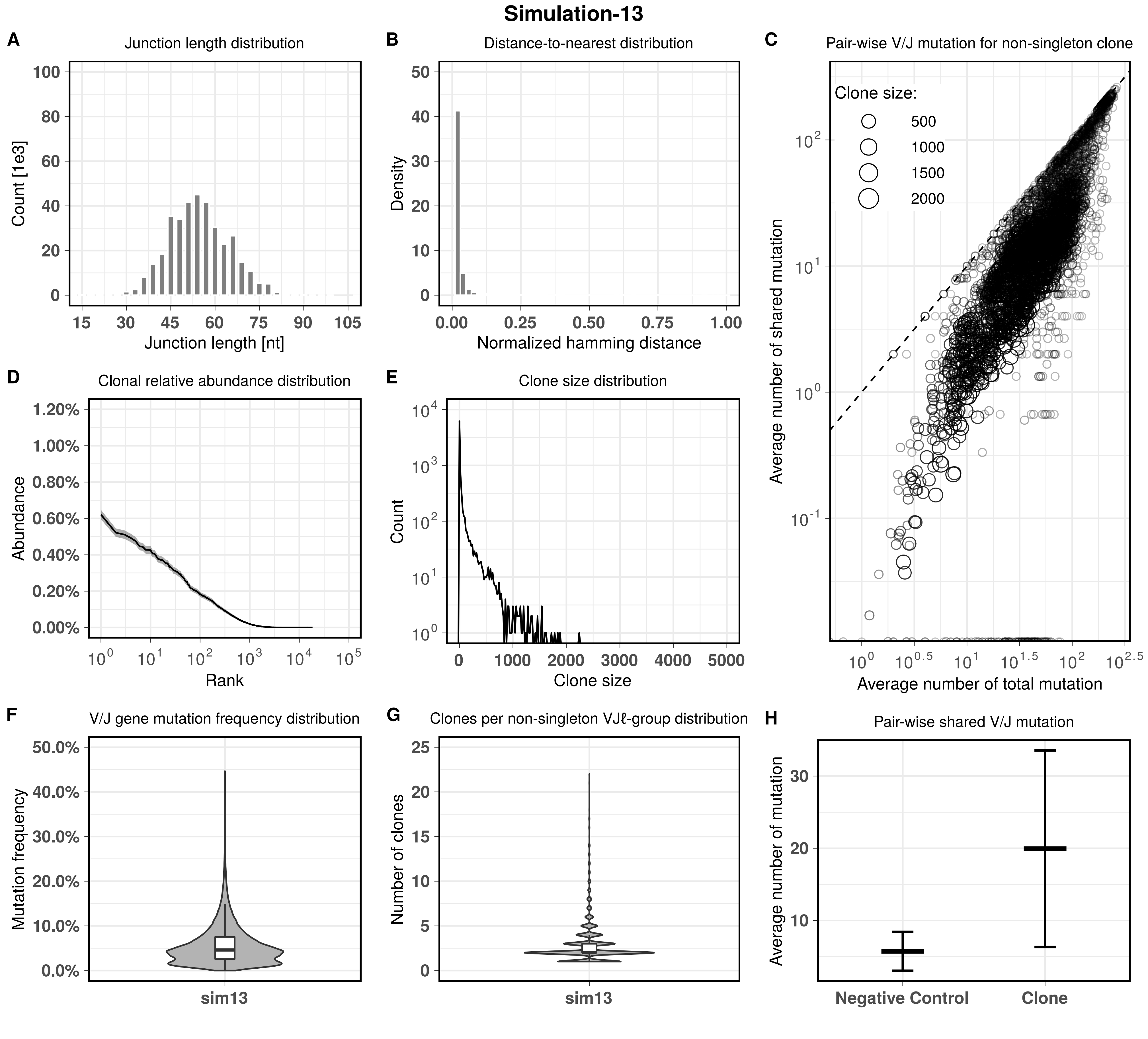

Simulation-13
A
Junction length distribution
Count [1e3]
Junction length [nt]
B
Distance-to-nearest distribution
Density
Normalized hamming distance
C
Pair-wise V/J mutation for non-singleton clone
Clone size:
500
1000
1500
2000
Average number of shared mutation
Average number of total mutation
D
Clonal relative abundance distribution
Abundance
Rank
E
Clone size distribution
Count
Clone size
F
V/J gene mutation frequency distribution
Mutation frequency
sim13
G
Clones per non-singleton VJℓ-group distribution
Number of clones
sim13
H
Pair-wise shared V/J mutation
Average number of mutation
Negative Control
Clone

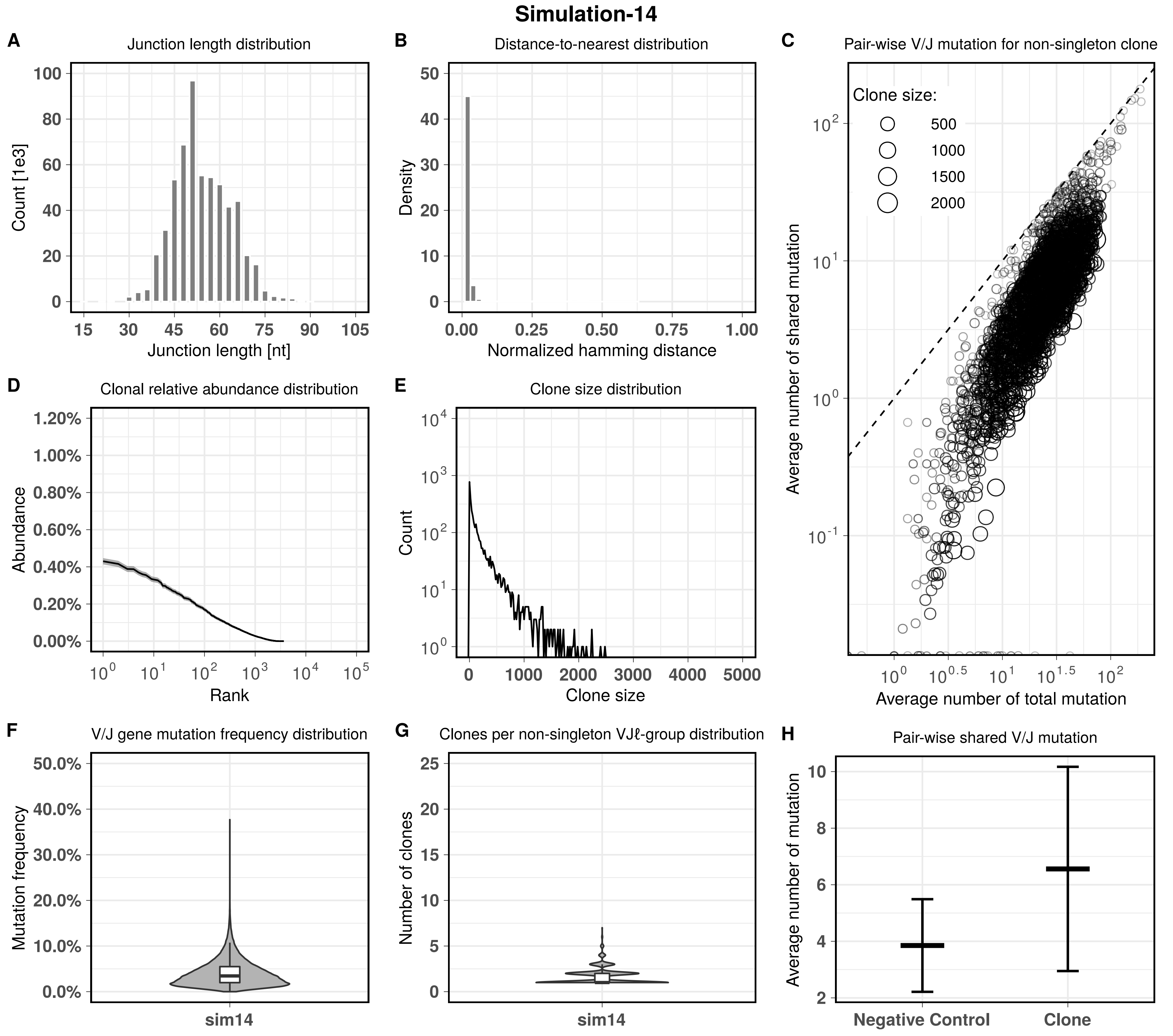

Simulation-14
A
Junction length distribution
Count [1e3]
Junction length [nt]
B
Distance-to-nearest distribution
Density
Normalized hamming distance
C
Pair-wise V/J mutation for non-singleton clone
Clone size:
500
1000
1500
2000
Average number of shared mutation
Average number of total mutation
D
Clonal relative abundance distribution
Abundance
Rank
E
Clone size distribution
Count
Clone size
F
V/J gene mutation frequency distribution
Mutation frequency
sim14
G
Clones per non-singleton VJℓ-group distribution
Number of clones
sim14
H
Pair-wise shared V/J mutation
Average number of mutation
Negative Control
Clone

### Simulation-15

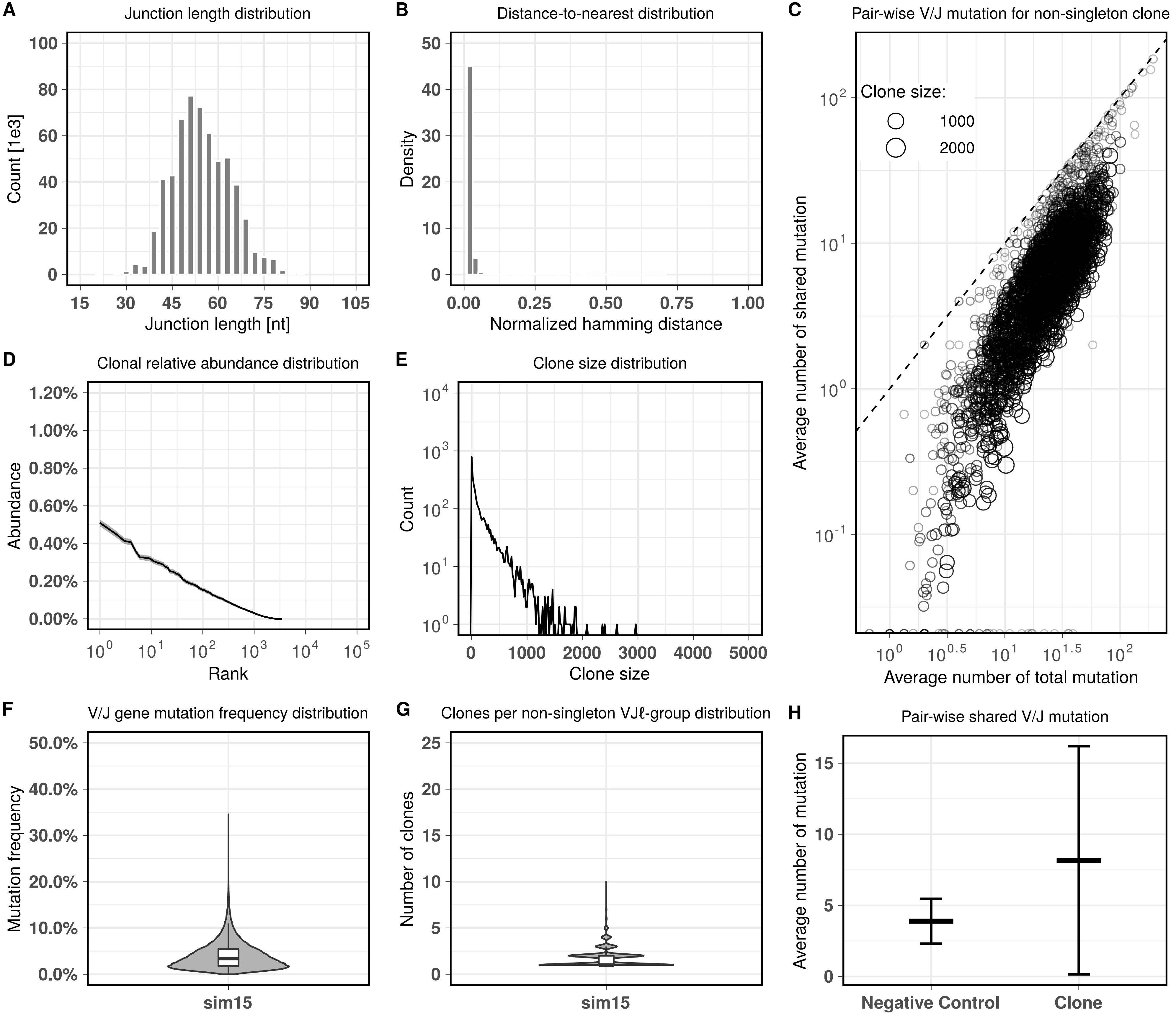

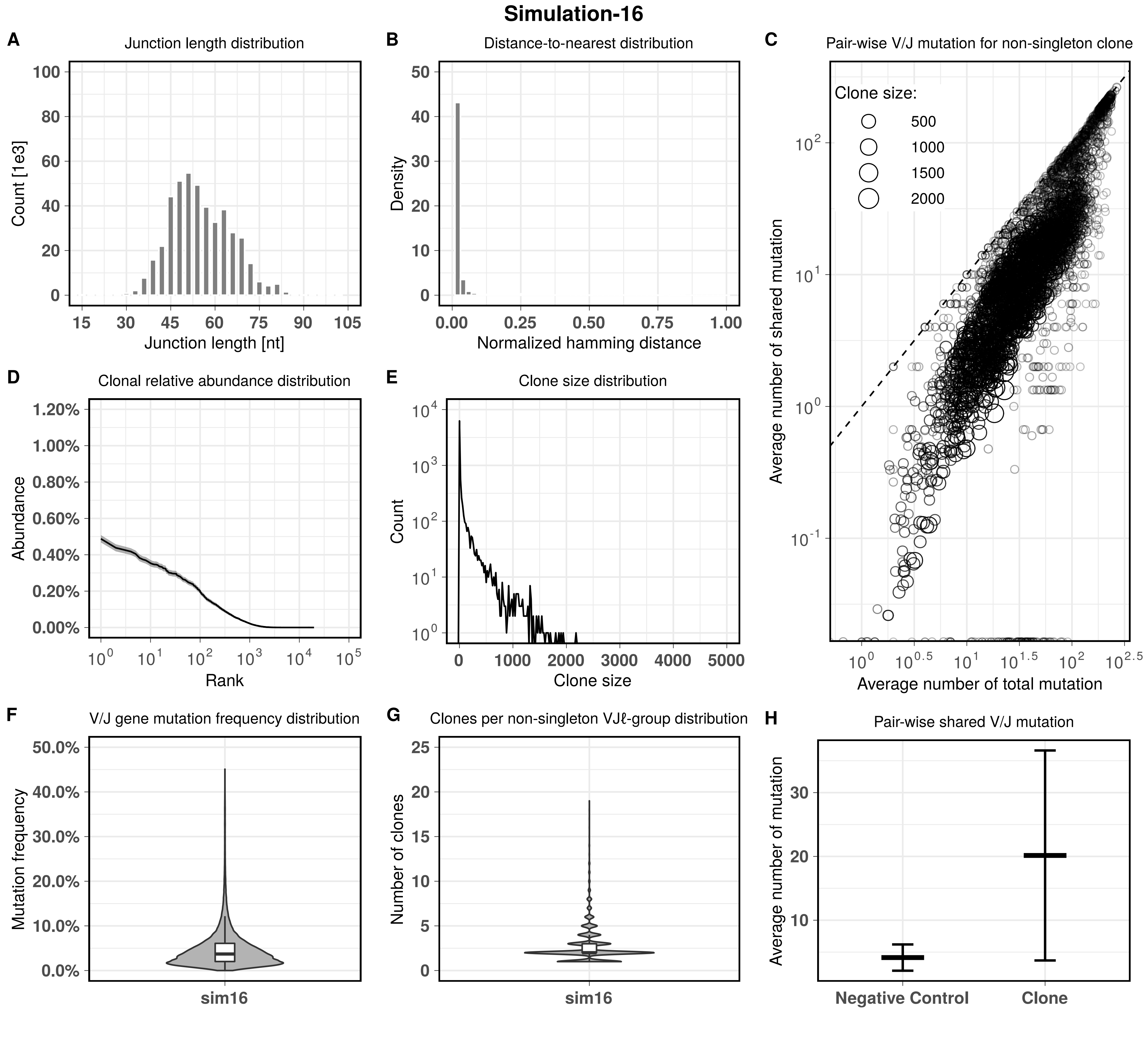
Simulation-16
A
Junction length distribution
Count [1e3]
Junction length [nt]
B
Distance-to-nearest distribution
Density
Normalized hamming distance
C
Pair-wise V/J mutation for non-singleton clone
Clone size:
500
1000
1500
2000
Average number of shared mutation
Average number of total mutation
D
Clonal relative abundance distribution
Abundance
Rank
E
Clone size distribution
Count
Clone size
F
V/J gene mutation frequency distribution
Mutation frequency
sim16
G
Clones per non-singleton VJℓ-group distribution
Number of clones
sim16
H
Pair-wise shared V/J mutation
Average number of mutation
Negative Control
Clone

#### Simulation-17

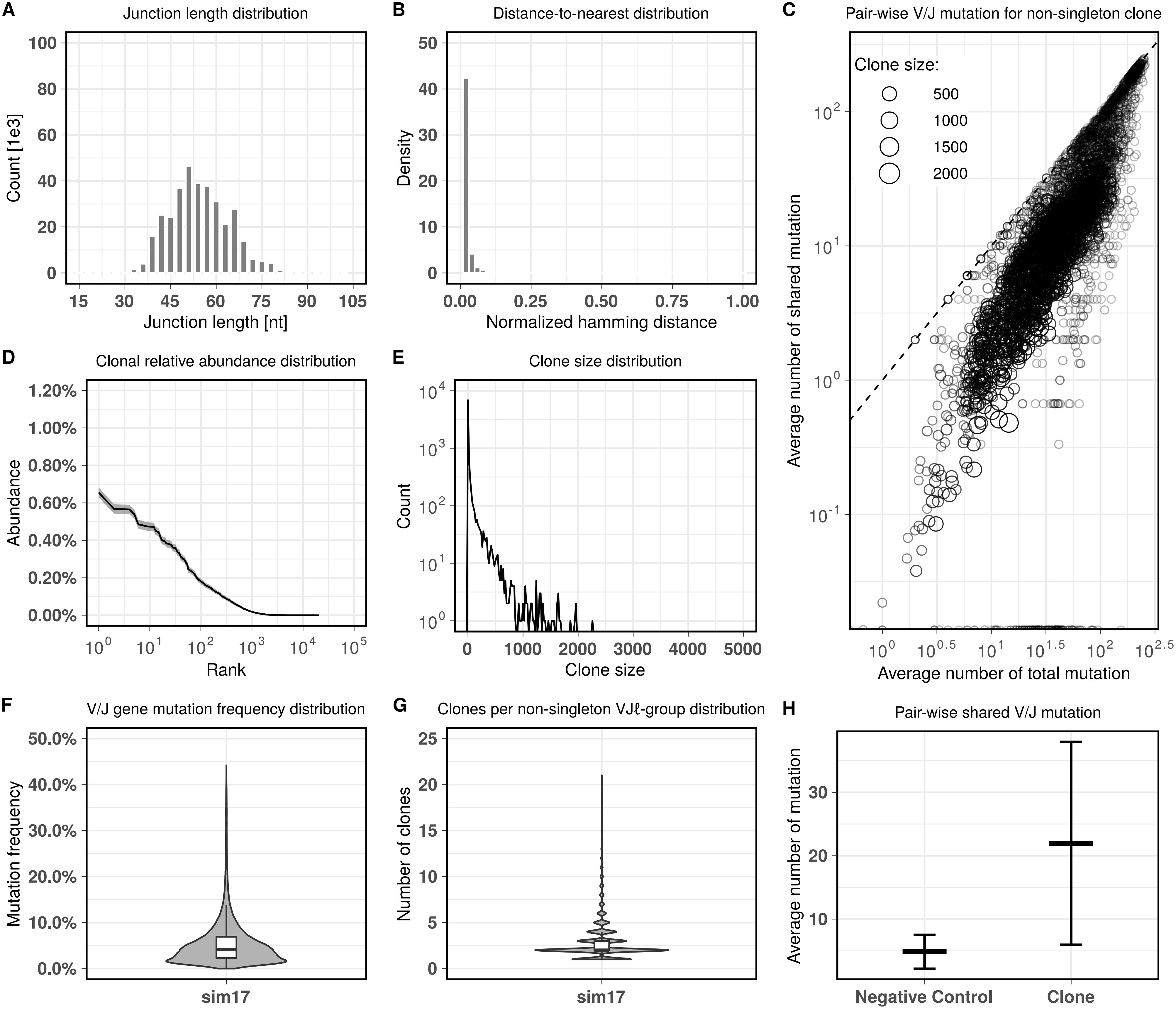

**Simulation-18**

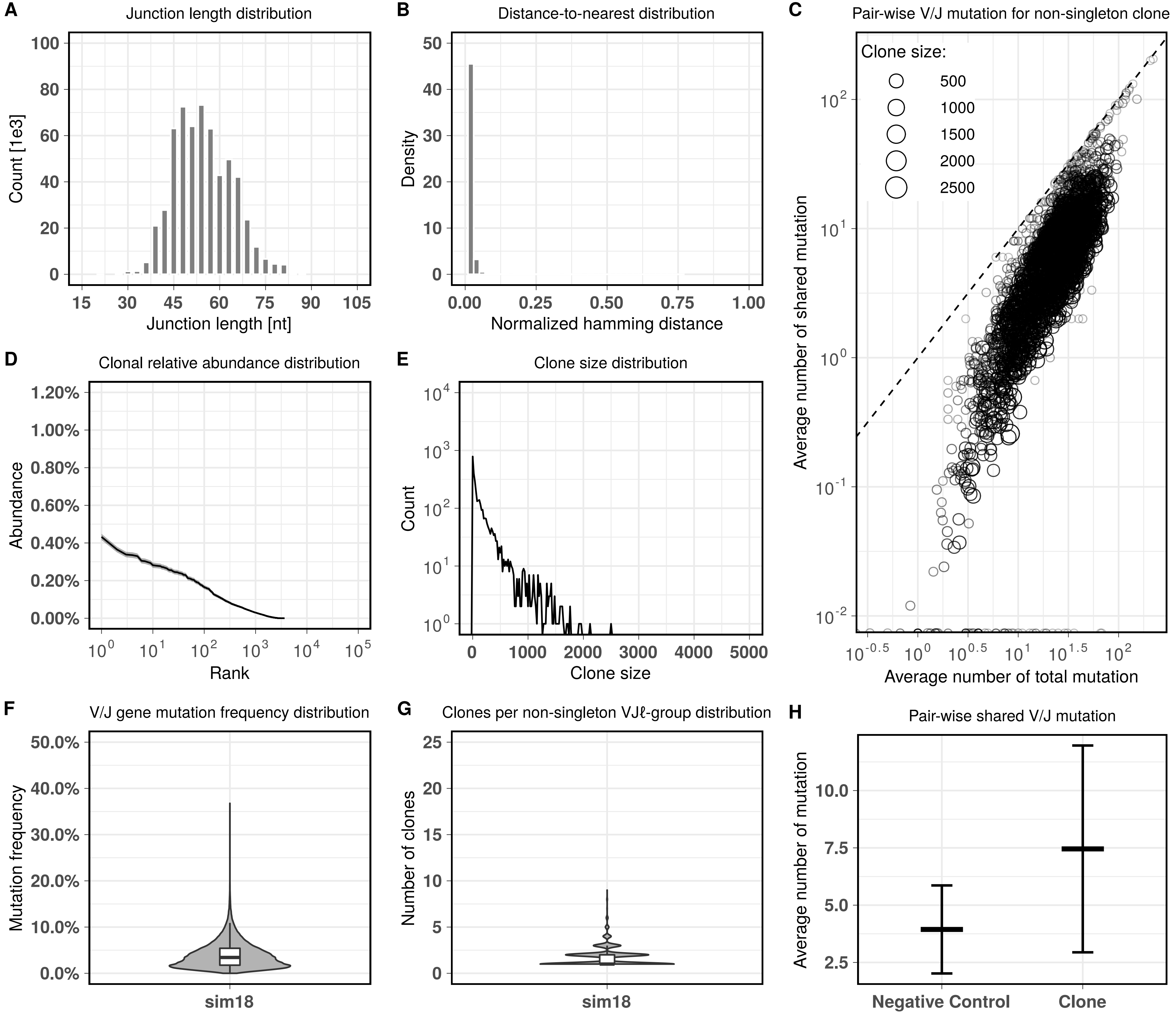

### Simulation-19

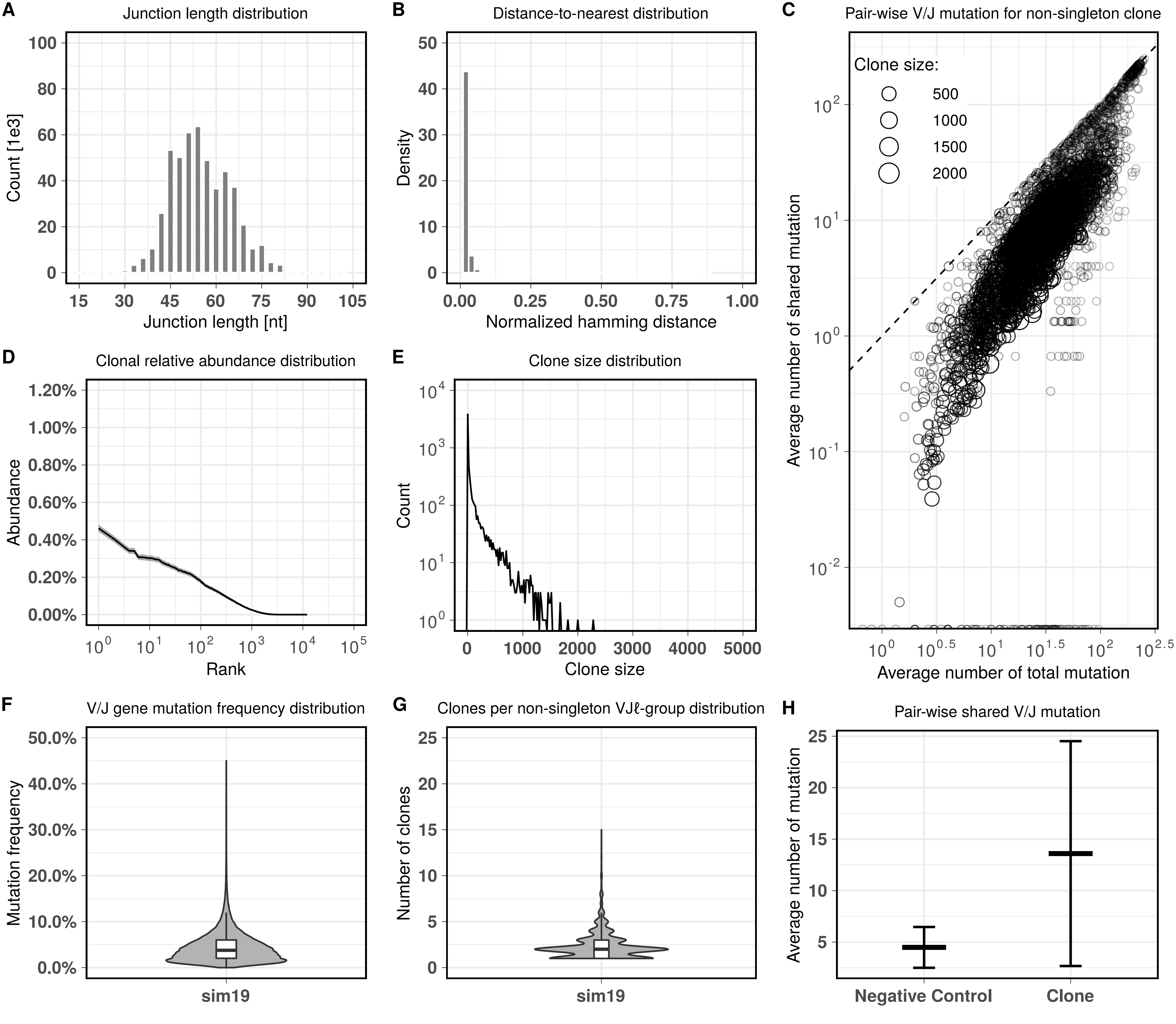

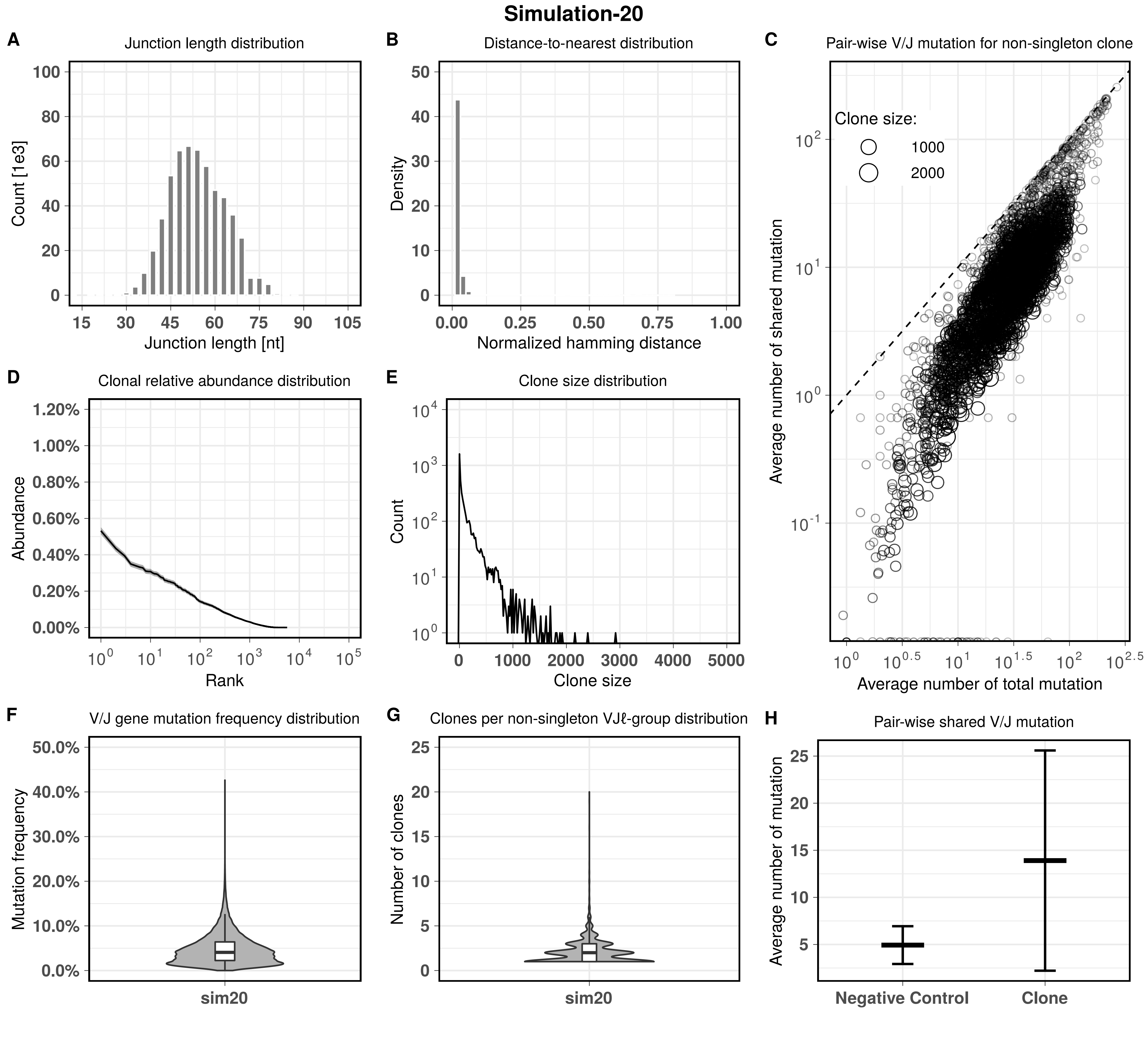
Simulation-20
A
Junction length distribution
Count [1e3]
Junction length [nt]
B
Distance-to-nearest distribution
Density
Normalized hamming distance
C
Pair-wise V/J mutation for non-singleton clone
Clone size:
1000
2000
Average number of shared mutation
Average number of total mutation
D
Clonal relative abundance distribution
Abundance
Rank
E
Clone size distribution
Count
Clone size
F
V/J gene mutation frequency distribution
Mutation frequency
sim20
G
Clones per non-singleton VJℓ-group distribution
Number of clones
sim20
H
Pair-wise shared V/J mutation
Average number of mutation
Negative Control
Clone

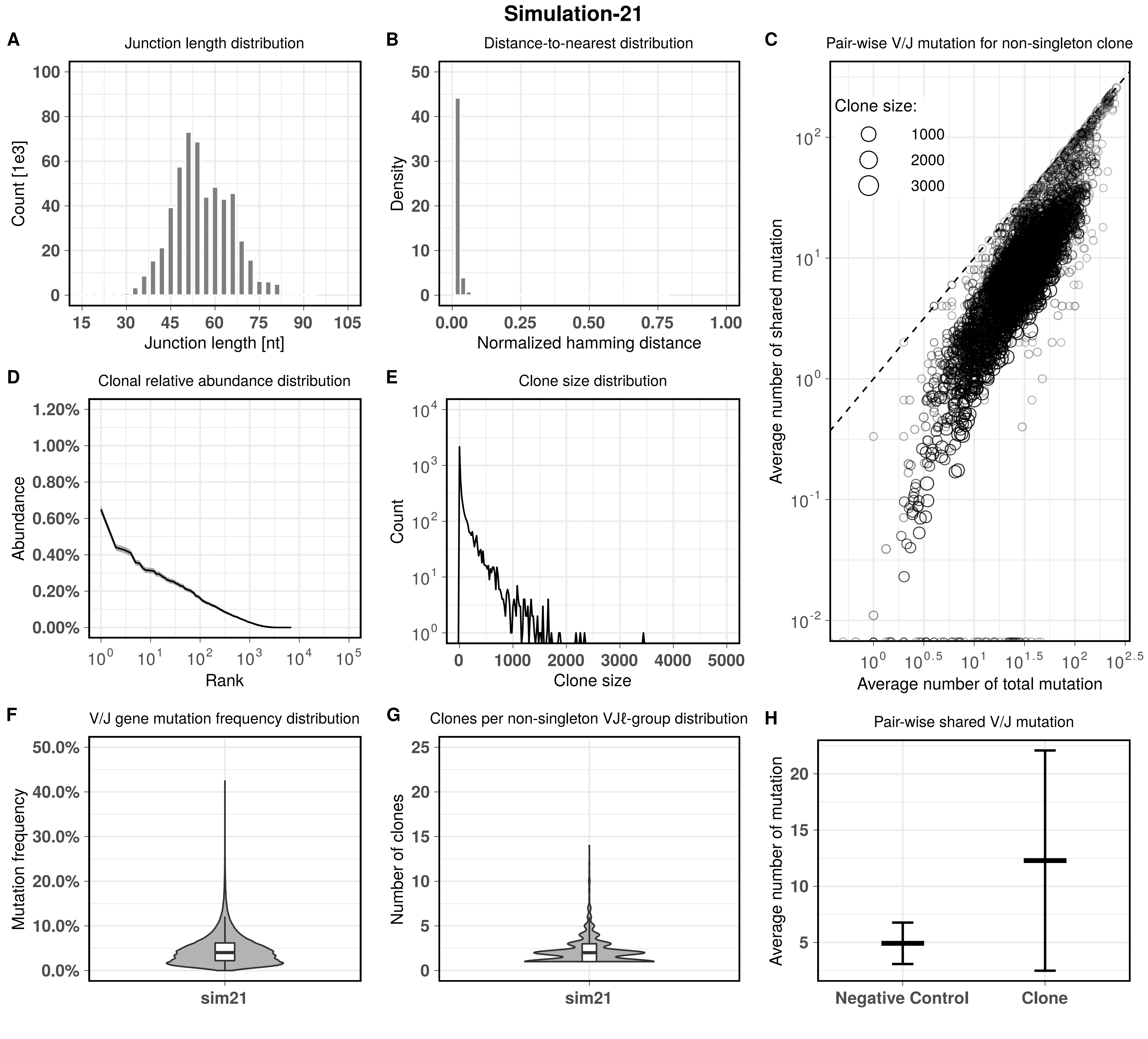

Simulation-21
A
Junction length distribution
Count [1e3]
Junction length [nt]
B
Distance-to-nearest distribution
Density
Normalized hamming distance
C
Pair-wise V/J mutation for non-singleton clone
Clone size:
1000
2000
3000
Average number of shared mutation
Average number of total mutation
D
Clonal relative abundance distribution
Abundance
Rank
E
Clone size distribution
Count
Clone size
F
V/J gene mutation frequency distribution
Mutation frequency
sim21
G
Clones per non-singleton VJℓ-group distribution
Number of clones
sim21
H
Pair-wise shared V/J mutation
Average number of mutation
Negative Control
Clone

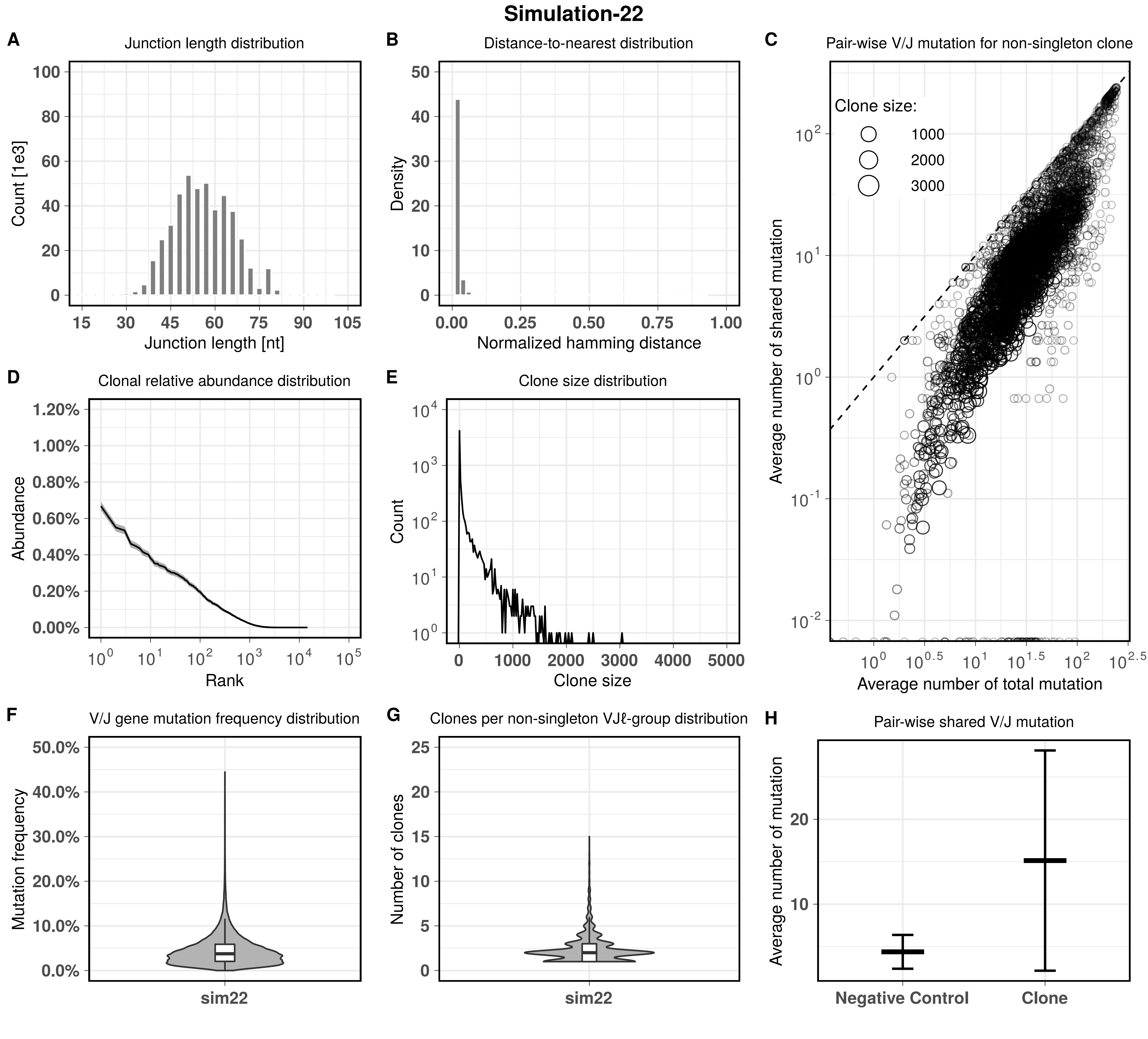
Simulation-22
A
Junction length distribution
Count [1e3]
Junction length [nt]
B
Distance-to-nearest distribution
Density
Normalized hamming distance
C
Pair-wise V/J mutation for non-singleton clone
Clone size:
1000
2000
3000
Average number of shared mutation
Average number of total mutation
D
Clonal relative abundance distribution
Abundance
Rank
E
Clone size distribution
Count
Clone size
F
V/J gene mutation frequency distribution
Mutation frequency
sim22
G
Clones per non-singleton VJℓ-group distribution
Number of clones
sim22
H
Pair-wise shared V/J mutation
Average number of mutation
Negative Control
Clone

### Simulation-23

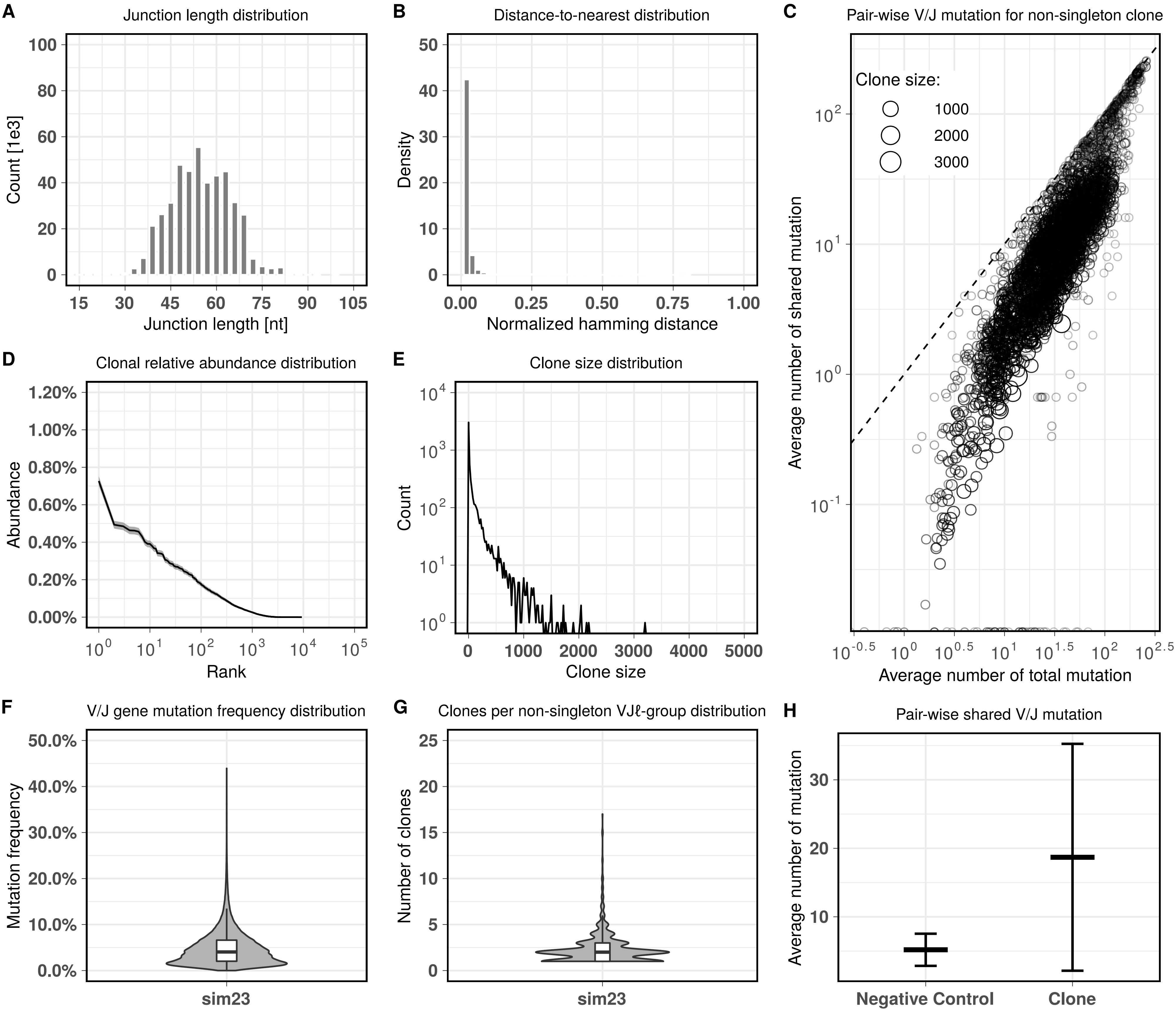

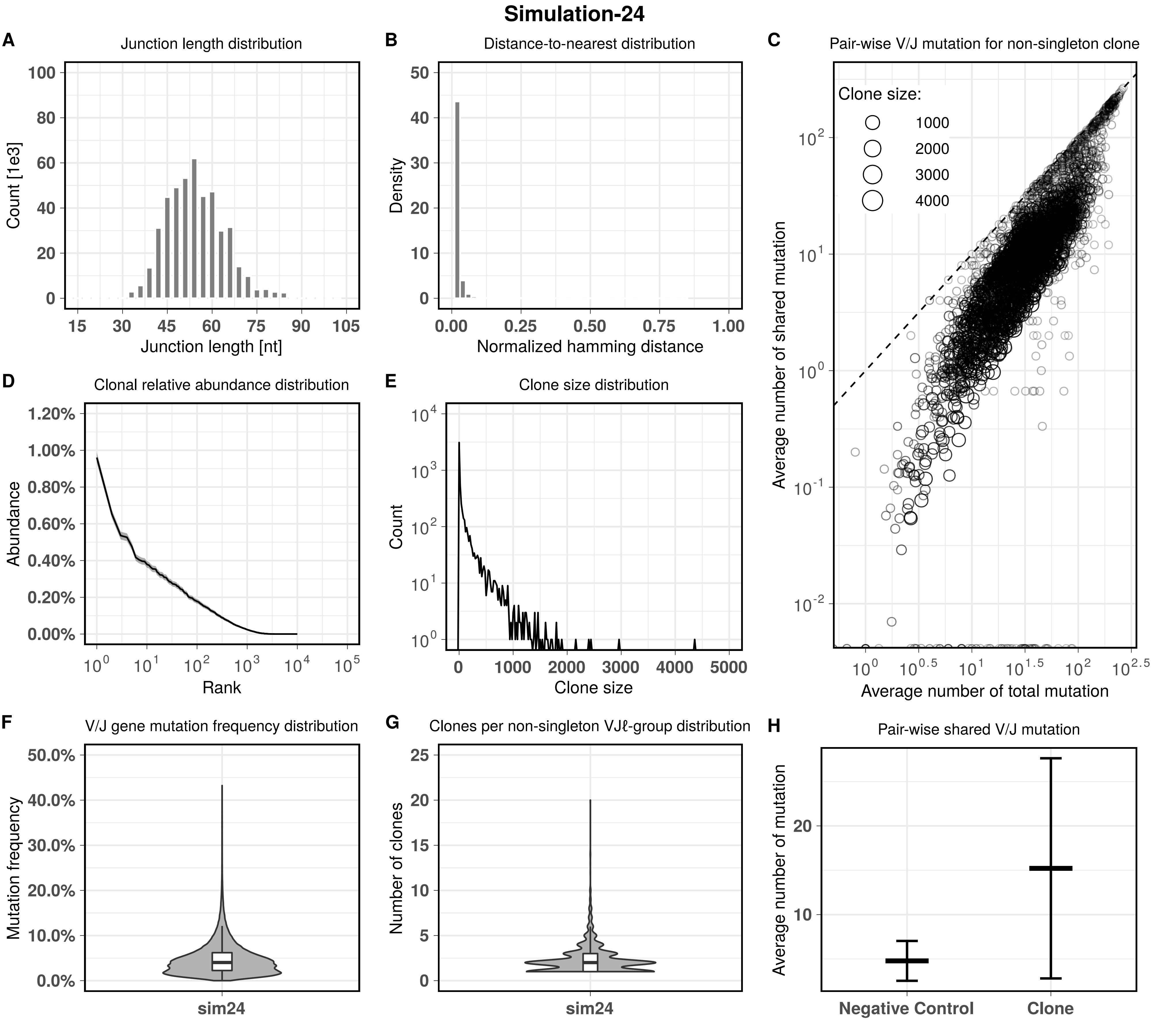
Simulation-24
A
Junction length distribution
Count [1e3]
Junction length [nt]
B
Distance-to-nearest distribution
Density
Normalized hamming distance
C
Pair-wise V/J mutation for non-singleton clone
Clone size:
1000
2000
3000
4000
Average number of shared mutation
Average number of total mutation
D
Clonal relative abundance distribution
Abundance
Rank
E
Clone size distribution
Count
Clone size
F
V/J gene mutation frequency distribution
Mutation frequency
sim24
G
Clones per non-singleton VJℓ-group distribution
Number of clones
sim24
H
Pair-wise shared V/J mutation
Average number of mutation
Negative Control
Clone

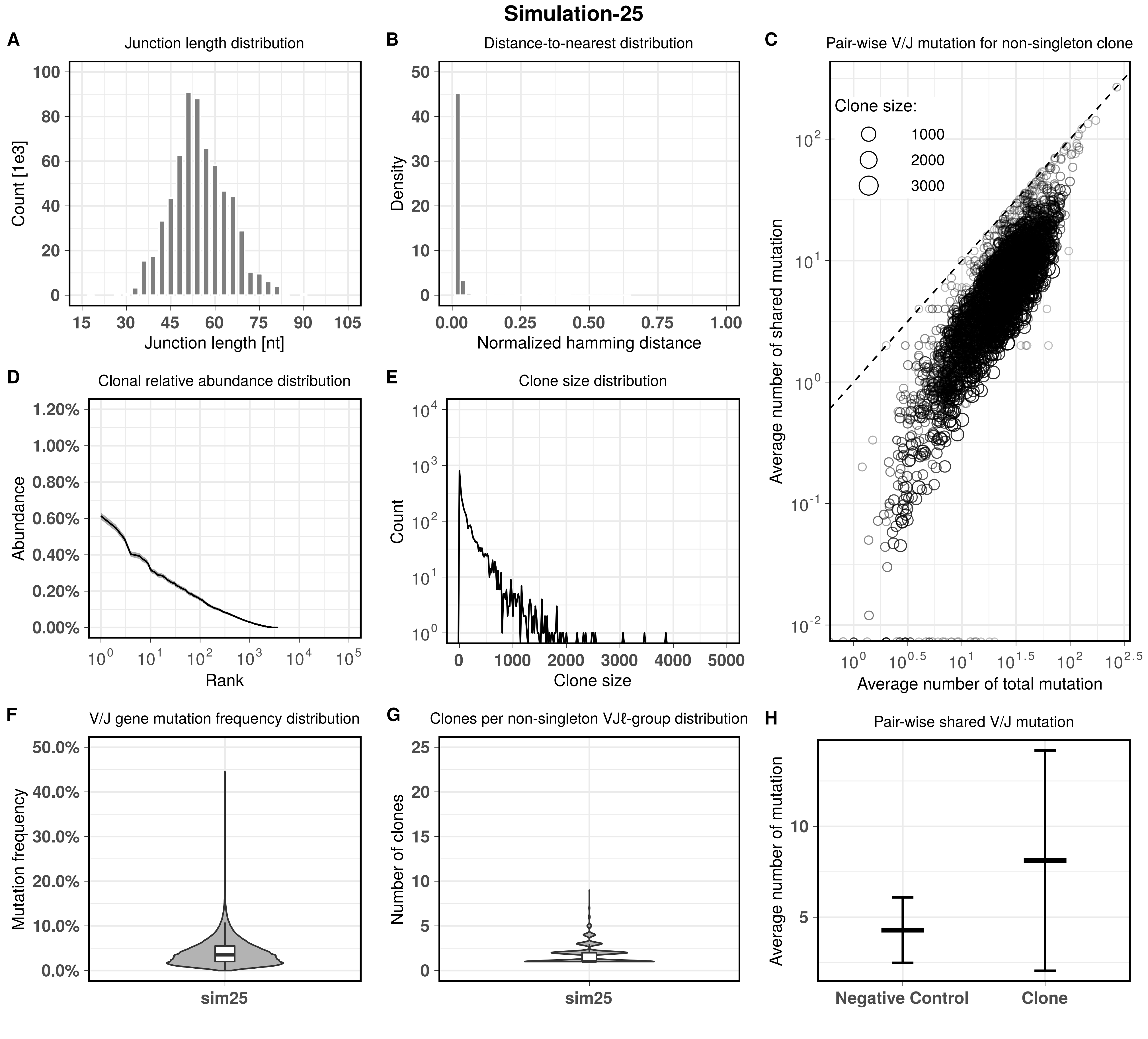

Simulation-25
A
Junction length distribution
Count [1e3]
Junction length [nt]
B
Distance-to-nearest distribution
Density
Normalized hamming distance
C
Pair-wise V/J mutation for non-singleton clone
Clone size:
1000
2000
3000
Average number of shared mutation
Average number of total mutation
D
Clonal relative abundance distribution
Abundance
Rank
E
Clone size distribution
Count
Clone size
F
V/J gene mutation frequency distribution
Mutation frequency
sim25
G
Clones per non-singleton VJℓ-group distribution
Number of clones
sim25
H
Pair-wise shared V/J mutation
Average number of mutation
Negative Control
Clone
